# Characterizing and targeting glioblastoma neuron-tumor networks with retrograde tracing

**DOI:** 10.1101/2024.03.18.585565

**Authors:** Svenja K. Tetzlaff, Ekin Reyhan, C. Peter Bengtson, Julian Schroers, Julia Wagner, Marc C. Schubert, Nikolas Layer, Maria C. Puschhof, Anton J. Faymonville, Nina Drewa, Rangel L. Pramatarov, Niklas Wissmann, Obada Alhalabi, Alina Heuer, Nirosan Sivapalan, Joaquín Campos, Berin Boztepe, Jonas G. Scheck, Giulia Villa, Manuel Schröter, Felix Sahm, Karin Forsberg-Nilsson, Michael O. Breckwoldt, Claudio Acuna, Bogdana Suchorska, Dieter Henrik Heiland, Julio Saez-Rodriguez, Varun Venkataramani

## Abstract

Glioblastomas are invasive brain tumors with high therapeutic resistance. Neuron-to-glioma synapses have been shown to promote glioblastoma progression. However, a characterization of tumor-connected neurons has been hampered by a lack of technologies. Here, we adapted retrograde tracing using rabies viruses to investigate and manipulate neuron-tumor networks. Glioblastoma rapidly integrated into neural circuits across the brain engaging in widespread functional communication, with acetylcholinergic neurons driving glioblastoma invasion. We uncovered patient-specific and tumor cell state-dependent differences in synaptogenic gene expression associated with neuron-tumor connectivity and subsequent invasivity. Importantly, radiotherapy enhanced neuron-tumor connectivity by increased neuronal activity. In turn, simultaneous neuronal activity inhibition and radiotherapy showed increased therapeutic effects, indicative of a role for neuron-to-glioma synapses in contributing to therapeutic resistance. Lastly, rabies-mediated genetic ablation of tumor-connected neurons halted glioblastoma progression, offering a viral strategy to tackle glioblastoma. Together, this study provides a framework to comprehensively characterize neuron-tumor networks and target glioblastoma.

## INTRODUCTION

Glioblastoma, the most prevalent and aggressive form of primary brain cancer in adults, presents a formidable challenge in neuro-oncology.^1,2^ Effective treatments remain elusive, largely due to the cellular heterogeneity, the highly invasive nature of glioblastoma and resistance to standard-of-care therapies including surgery, radio- and chemotherapy.^1,3–11^ A burgeoning area of interest is the exploration of the intricate relationships between glioblastoma cells and neural networks of the brain.^12–15^ The interplay between tumor cells and neuronal circuits, particularly synaptic neuron-tumor communication, has emerged as a critical factor in tumor progression and invasion.^12,16–25^ However, while neuronal molecular signatures have been described in paired primary and recurrent glioblastoma^4,7^, it is unclear whether and how neuron-glioma synaptic communication contributes to therapeutic resistance. Synaptic inputs onto adult glioblastoma cells have so far been identified as local, glutamatergic projections, leaving the comprehensive circuit architecture and the diversity of neuronal subtypes interacting with glioma largely unexplored.^8,16–18^ Moreover, the dynamics of how tumor cells synaptically integrate into neuronal networks and in turn change neuronal structure and function are yet unclear. The cellular, molecular and functional heterogeneity of glioblastoma has been increasingly investigated,^5,8–10,26^ but how these layers are related to neuronal connectivity is yet unknown.

While tracing neuronal circuits is an extensive field of research in neuroscience,^27–29^ the neuronal connectome of brain tumors remains poorly understood.^12,13^ Among tracing approaches, the retrograde monosynaptic tracing using modified rabies virus stands out as a pivotal technique for investigating neural networks.^30–33^ Previous studies have applied this methodology to neurons and oligodendrocytic precursor cells, both receiving synaptic input,^34,35^ to map their neuronal connectome and characterize their functional organization.^33,36–44^

This paper introduces a modified rabies virus-based retrograde tracing methodology platform for the multimodal, neuronal connectome characterization of glioblastoma. We demonstrated its applicability across model systems ranging from human patient tissue, patient-derived xenograft models to co-cultures of neurons and tumor cells. Unexpectedly, this approach revealed a majority of glioblastoma cells and neurons were functionally connected to neurons in the early stages of glioblastoma colonization. This stands in contrast to previous data from us and others,^16,17^ where technologies to comprehensively assess the functional connectivity were lacking. Molecular and functional analyses of tumor-connected (connected^TUM^) and tumor-unconnected (unconnected^TUM^) neurons did not show significant differences in early stages of colonization, implying that synaptic integration of tumor cells into neural circuits precedes neuronal dysfunction and hyperexcitability, described in later stages of the disease.^45–49^ Moreover, we found brain-wide recruitment of diverse neuronal populations including neuromodulatory circuits forming neuron-tumor networks with glioblastoma. Together, both acetylcholinergic and glutamatergic neurons were able to drive glioblastoma progression. Further, invasive patient-derived tumors and glioblastoma cell states were associated synaptogenic gene expression signatures and subsequent larger neuron-tumor connectivity. Lastly, we found that radiotherapy promotes neuron-tumor connectivity by boosting neuronal activity and saw an increased therapeutic effect of combined neuronal activity inhibition and radiotherapy. Hereby, we provided evidence for the role of neuron-to-glioma synaptic communication in contributing to therapeutic resistance. Lastly, we provided a proof-of-concept of how, in addition to pharmacological perturbation, rabies virus itself could be used to selectively ablate connected^TUM^ neurons and thereby inhibit glioblastoma progression.

The insights gathered here offer a valuable framework for future investigations in glioblastoma and potentially other cancer entities, highlighting the pivotal role of characterizing the neuronal connectome of glioblastoma to develop novel therapeutic strategies.

## RESULTS

### Rabies-based retrograde tracing enables versatile neuron-tumor network characterization

We took advantage of a rabies virus-based retrograde tracing system to establish a method for characterizing neuron-tumor networks using patient-derived glioblastoma spheroid cultures (Figures 1A, B).^8,9,50,51^ First, we stably transduced glioblastoma spheroids (n = 10 patient-derived models, Figure 1B, Supplementary Table 1) with a lentivirus containing the EnvA receptor TVA for rabies entry, the rabies virus glycoprotein (oG) for trans-complementation as well as spread, and the cytosolically expressed fluorophore mCherry (STAR Methods). Second, we performed fluorescence-associated cell sorting (FACS) for mCherry, to identify and isolate patient spheroid cells expressing TVA and oG. Subsequently, these cells were transduced with an EnvA-pseudotyped G protein-deleted (ΔG) rabies virus expressing the cytosolic fluorophore GFP that could only infect glioblastoma cells containing the TVA receptor. Upon entry and trans-complementation with the rabies-oG protein, starter glioblastoma cells (GB^Starter^) are expected to label tumor cell-connected (connected^TUM^) neurons via monosynaptic, retrograde propagation.^31^ Connected^TUM^ neurons could be readily identified by expressing only GFP, whereas patient-derived GB^Starter^ cells expressed both mCherry and GFP. Further, as connected^TUM^ neurons did not express oG, no transmission across secondary synapses was possible, ensuring a high specificity of this approach to label only directly connected neuron-tumor networks (Figure 1A). FACS of GB^Starter^ spheroids enabled a direct and dense labeling of all tumor cells before engrafting these tumor cells into any experimental model system. In contrast, implanting or seeding glioblastoma cells before (ΔG) rabies virus transduction in combination with a titration of the ΔG rabies virus (STAR Methods) led to a sequential, sparse labeling to trace the neuronal connectome of single glioblastoma cells. While whole-tumor neuronal connectome analyses enabled a comprehensive characterization, sparse labeling approaches allowed for a more specific analysis of single tumor cell states and their connected^TUM^ neurons over time. Further, genetic modification of the rabies virus additionally expressing functional proteins such as the Cre Recombinase^52^ enabled a precise manipulation of connected^TUM^ neurons to investigate its subsequent effect on glioblastoma biology (Figures 1A, S1A).

**Figure 1.**
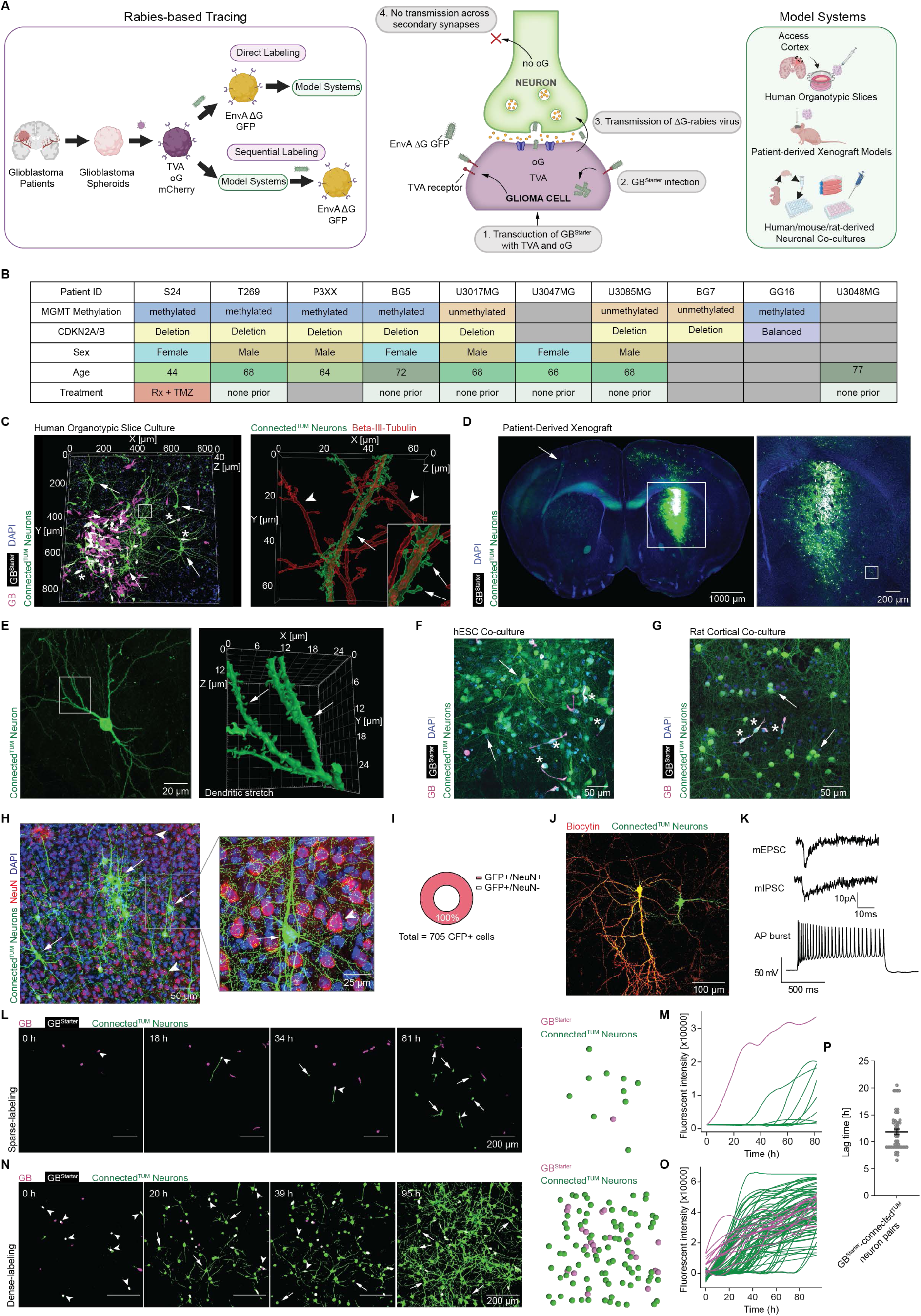
Rabies-based tracing of glioblastoma neuron-tumor networks across model systems. (A) Monosynaptic retrograde tracing workflow in patient-derived glioblastoma (GB) spheroid models. (B) Overview of patient-derived glioblastoma models used in this study. (C) 3D rendering of an exemplary human organotypic slice injected with S24 GB^Starter^ cells (left). Shown are GB^Starter^ cells (white, asterisks) and connected^TUM^ neurons (CVS-N2c^ΔG^-eGFP(EnvA), green, arrows) nearby. The inset (right) shows the neuronal marker beta-III-tubulin (red) expressing dendritic stretch of a connected^TUM^ neuron (green, arrow). Arrowheads point to only beta-III-tubulin expressing, unconnected^TUM^ neurons (red). Zoom-in showing dendritic spines (arrows) of a connected^TUM^ neuron. (D) Retrograde tracing in a patient-derived xenograft model (PDX). Shown is an exemplary brain section, where S24 GB^Starter^ cells (white) and connected^TUM^ neurons (SAD-B19^ΔG^-eGFP(EnvA), green) are visible. Arrow points to distant connected^TUM^ neurons on the contralateral hemisphere. The inset is a zoom-in on the tumor site (dashed white circle). (E) An exemplary connected^TUM^ neuron is shown from the brain slice from D (left). 3D rendering of dendritic stretches (right) showing that dendritic spines (arrows) can be distinguished in connected^TUM^ neurons. (F) Retrograde tracing in human embryonic stem cell-induced neurons with human S24 GB^Starter^ cells (white). Arrows show exemplary connected^TUM^ neurons (CVS-N2c^ΔG^-eGFP(EnvA), green). Asterisks point to GB^Starter^ cells. (G) Retrograde tracing in co-culture of rat cortical neurons with human S24 GB^Starter^ cells (white). Arrows show exemplary connected^TUM^ neurons (CVS-N2c^ΔG^-eGFP(EnvA), green). Asterisks point to GB^Starter^ cells. Confocal imaging of connected^TUM^ neurons (CVS-N2c^ΔG^-eGFP(EnvA), green) in a tissue section from PDX model S24 stained against the neuronal marker NeuN (red). Arrows point to connected^TUM^ neurons and arrowheads show only NeuN-positive, unconnected^TUM^ neurons. (H) Quantification showing the portion of NeuN-positive cells from the retrogradely labeled, connected^TUM^ neurons (SAD-B19^ΔG^-eGFP(EnvA), green) (n = 705 GFP-positive cells in n = 10 different patient-derived GB models). (I) Confocal imaging of a patched connected^TUM^ neuron (CVS-N2c^ΔG^-eGFP(EnvA), green) filled with Neurobiotin and stained for streptavidin 647 (yellow). (J) Representative examples of mEPSC (top), mIPSC (middle) and AP bursts after current injection (bottom) of a connected^TUM^ neuron by whole-cell patch clamp recording. (K) Probability maps of live cell time-lapse imaging demonstrating the sparse labeling approach. One of many S24 GB cells (magenta) is labeled with rabies virus and becomes a GB^Starter^ cell (white, arrowheads). Over a time course of 81h the neuronal connectome of this GB^Starter^ cell is traced and connected^TUM^ neurons become infected (CVS-N2c^ΔG^-eGFP(EnvA), green, arrows). Rendered manual segmentation representing the last time point of imaging (far right). Each dot represents a cell. (L) Line plot indicating the change of (CVS-N2c^ΔG^) eGFP fluorescence intensities of the GB^Starter^ cell (magenta line) and its connected^TUM^ neurons (green lines) as shown in L. (CVS-N2c^ΔG^) eGFP fluorescence is indicative of rabies virus infection. (M) Probability maps of live cell time-lapse imaging showing the dense labeling approach. Most GB cells are labeled with rabies virus and become double-positive (white, arrowheads). Over 95h the neuronal connectome of the whole tumor region imaged is traced. Arrows point to exemplary connected^TUM^ neurons. Rendered manual segmentation representing the last time point of imaging (far right). Each dot represents a cell. (N) Line plot indicating the change of the (CVS-N2c^ΔG^) eGFP fluorescence intensities of the GB^Starter^ cells (magenta lines) and their connected^TUM^ neurons (green lines) as shown in N. (CVS-N2c^ΔG^) eGFP fluorescence is indicative of rabies virus infection. (O) Quantification of the lag time with which (SAD-B19^ΔG^/CVS-N2c^ΔG^) eGFP fluorescence can be observed in connected^TUM^ neurons after their respective GB^Starter^ cells have been infected with rabies via GB^Starter^ cells (n = 49 GB^Starter^-connected^TUM^ neuron pairs analyzed).

We assessed the versatility of this approach for tracing of functional neuron-tumor networks across a range of *in vivo, ex vivo* and *in vitro* model systems. To establish an all-in-human tissue model system for tracing neuron-tumor networks, we adapted an organotypic slice culture using human access cortex tissue removed during surgery^53^ (n = 7 patients, Figure 1C) and transplanted GB^Starter^ cells to label human connected^TUM^ neurons. This model system was complemented by patient-derived mouse xenografts and a variety of human and mouse neuronal co-culture models (STAR Methods) (Figures 1C-G, S1B, C).^54^

Specifically, our retrograde tracing technique selectively labeled connected^TUM^ neurons as demonstrated across all patient-derived spheroids in all model systems and using different strains of the rabies virus (CVS-N2c^ΔG^-eGFP(EnvA)^33^ and SAD-B19^ΔG^-eGFP(EnvA)^30^ (Figures 1C-G). This labeling approach even allowed the ultrastructural characterization of connected^TUM^ neurons including different classes of dendritic spines^55^ employing high- and super-resolution light microscopy (Figures 1C, E, S1D). Our analysis confirmed that retrogradely labeled cells are exclusively neuronal, with no labeling observed in astrocytes, oligodendrocytes, or microglia (Figures 1H, I, S1E-G). Electrophysiological assessments revealed connected^TUM^ neurons maintained their characteristic functional properties, including action potential firing, bursting as well as excitatory and inhibitory synaptic inputs from other neurons (Figures 1J, K, S1H). Importantly, our tracing method was highly specific, as close to no labeling occurred in the absence of TVA receptor expression in glioma cells, nor when media from neuron-GB^Starter^ co-cultures was added to untransduced neuronal cultures (Figures S1I, J).

Last, we evaluated the tumor cell-toxic potential of two strains of the rabies virus, CVS-N2c^ΔG^-eGFP(EnvA) and SAD-B19^ΔG^-eGFP(EnvA).^30,33^ Here, we found little sign of tumor cell toxicity, as tumor cells transduced with either of these rabies strains showed comparable growth curves in monocultures as compared to control cell lines (Figure S1K).

In summary, rabies-based retrograde tracing in glioblastoma enables a comprehensive investigation of connected^TUM^ neurons across a variety of model systems.

### Rapid and dynamic integration of glioblastoma into neuron-tumor networks

Live-cell imaging of neuron-tumor network formation in co-culture models over time revealed the fast and dynamically increasing integration of glioblastoma cells into neuronal circuits. Sparse labeling of GB^Starter^ cells enabled tracking the recruitment of connected^TUM^ neurons in a near real-time manner, increasing over time (Figures 1 L, M, Video S1). In contrast, employing dense labeling to mark the entirety of the tumor cell population permits a comprehensive examination of the neuronal connectome associated with all tumor cells. This approach contrasts with the sparse labeling technique, which reveals neuronal connections to individual tumor cells, by enabling the visualization of neuronal networks linked to the entire tumor (Figures 1N, O, Video S1). Remarkably, connected^TUM^ neuron labeling occurred within a matter of hours (mean 11.85 +/- 0.51 hours) after GB^Starter^ cells became GFP-positive, demonstrating the rapid ability of glioblastoma cells to form neuron-tumor connections, in contrast to a previously reported minimum amount of two days to retrogradely label neuron-to-neuron synapses (Figures 1P, S1L).^31^ These findings were complemented by corresponding electrophysiological measurements of neuronal activity-driven excitatory, postsynaptic currents and slow inward currents (Figure S1M).^8,16–18^

Together, combined retrograde tracing and live-cell imaging revealed a rapid, functional integration of glioblastoma cells into neuronal circuits.

### Widespread functional neuron-tumor network communication in glioblastoma

We aimed to understand the structural and functional connectivity between neurons linked to tumor cells. Unexpectedly, we found in the early stages of glioblastoma colonization, all tumor clusters label connected^TUM^ neurons, indicative of a high level of structural connectivity between neurons and glioblastoma cells (Figure 2A). To characterize whether these are corresponding to functional neuron-tumor networks, we performed paired whole-cell patch-clamp electrophysiology of putatively connected^TUM^ neurons and glioblastoma cells in co-cultures of patient-derived glioblastoma cells and neurons (Figure 2B). This allowed us to examine their electrophysiological and functional connectivity (Figures 2C-E). We found that action potentials of connected^TUM^ neurons correlate with either excitatory postsynaptic currents (EPSCs) or slow inward currents (SICs) in glioma cells (Figures 2F, G).^16,17^ These co-active electrical activity patterns indicate robust functional connectivity between connected^TUM^, GFP-positive neurons and their corresponding GB^Starter^ cells.

**Figure 2.**
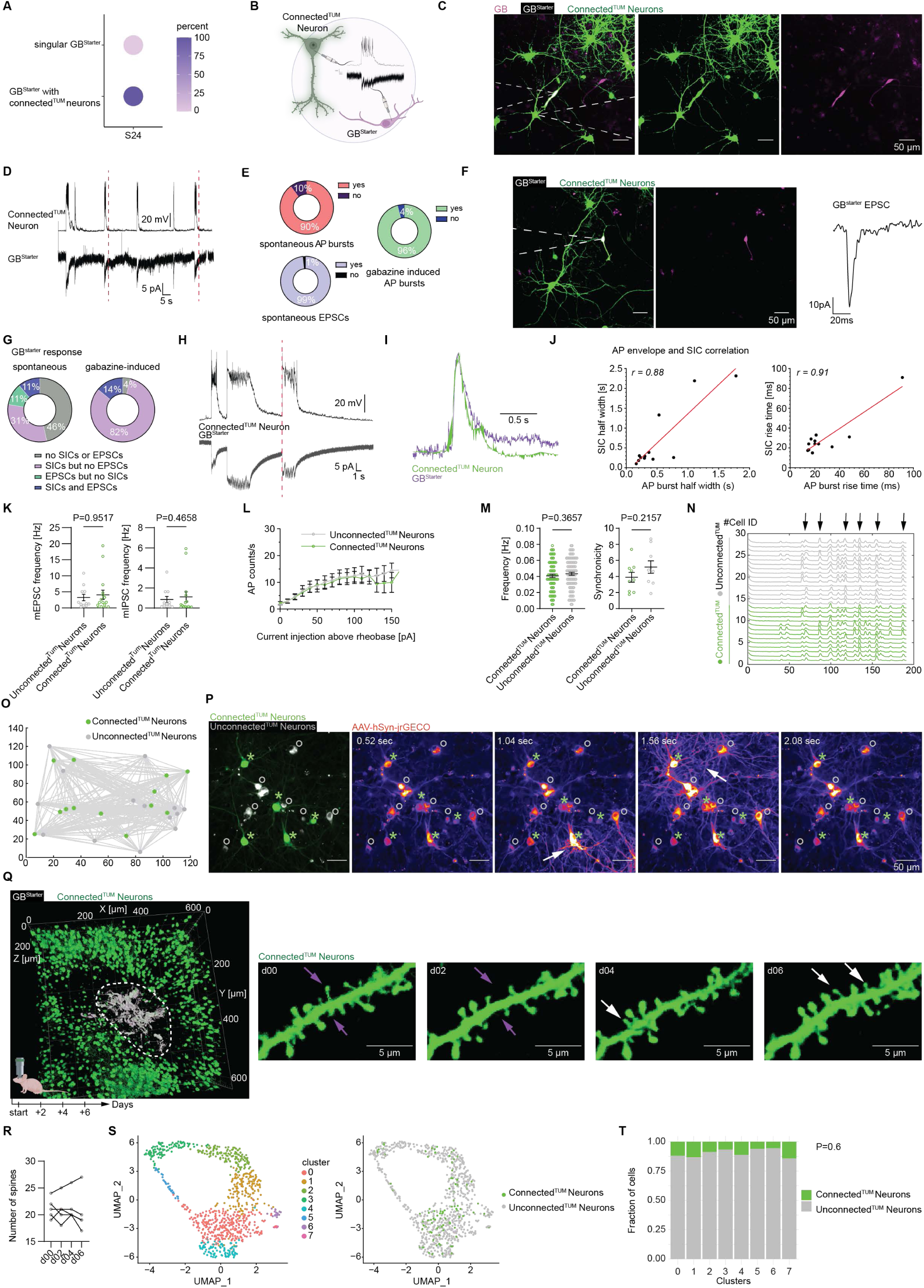
Functional investigation of neuron-tumor networks. (A) Quantification of GB^Starter^ cell connectivity percentage in co-cultures (n = 2529 GB^Starter^ cells in 10 samples). (B) Schematic of paired whole-cell patch clamp electrophysiology of connected^TUM^ neurons and GB^Starter^ cells. (C) Representative image of a S24 GB^Starter^ cell (white) and adjacent connected^TUM^ neuron in co-culture (CVS-N2c^ΔG^-eGFP(EnvA), green). Dashed white lines indicate patch pipettes of paired patch. (D) Exemplary electrophysiological traces of a connected^TUM^ neuron (top) and its respective S24 GB^Starter^ cell (bottom). Red dashed lines indicating synchronized events. (E) Quantification of neuronal activity of connected^TUM^ neurons with spontaneous AP bursts (left, top, n = 59 cells), spontaneous EPSCs (left, bottom, n = 59 cells) and gabazine-induced AP bursts (right, n = 25 cells). (F) Representative image of patched S24 GB^Starter^ cell (white, CVS-N2c^ΔG^-eGFP(EnvA)) and corresponding EPSC trace. Dashed white lines indicate patch pipette. (G) Quantification of electrophysiological GB^Starter^ response in the form of no response, only EPSCs, only SICs or both, under baseline condition (left, n = 63 pairs) and after stimulation with gabazine (right, n = 28 pairs). (H) Representative traces of paired-patched connected^TUM^ neuron (top) and GB^Starter^ cell (bottom) showing neuronal AP bursts and responsive SICs. Synchronized electrophysiological traces indicated by red dashed line. (I) Exemplary overlay of AP burst slope and GB^Starter^ cell SIC. (J) Correlation of AP envelopes and SICs. SIC halfwidth and AP burst half width (left), n = 12 pairs, Pearson’s r = 0.88, ANOVA F (df) = 33.8 (11), p = 0.0017. SIC rise time and AP burst rise time (right), Pearson’s r = 0.91, ANOVA F (df) = 49.8 (11), p = 0.00035. (K) Mean frequencies of mEPSCs and mIPSCs (n = 11 unconnected^TUM^ and n = 15-16 connected^TUM^ neurons, Mann-Whitney test). (L) Input-output relationship between the current injected relative to the rheobase current and the number of action potentials generated over 1 s in connected^TUM^ (n = 10) and unconnected^TUM^ (n = 9) regular-spiking neurons. (M) Calcium transient frequency (left) and synchronicity (right) of connected^TUM^ and unconnected^TUM^ neurons (n = 75 connected^TUM^ and 95 unconnected^TUM^ neurons in 9 regions of interest, Mann-Whitney test (frequency) and unpaired t-test (synchronicity). (N) Representative individual calcium traces of connected^TUM^ and unconnected^TUM^ neurons. Arrows pointing to exemplary synchronized events affecting all neurons. (O) Exemplary connected^TUM^ and unconnected^TUM^ calcium coactivity map. (P) Dual-color calcium imaging of unconnected^TUM^ (gray) and connected^TUM^ (CVS-N2c^ΔG^-eGFP(EnvA), green) neurons using AAV-jrGECO (fire) in co-culture. Asterisks show connected^TUM^ neurons, circles point to unconnected^TUM^ neurons. (Q) 3D rendering of *in vivo* two-photon longitudinal imaging of S24 GB^Starter^ cells (white) and connected^TUM^ neurons (CVS-N2c^ΔG^-eGFP(EnvA), green). Tumor overview is shown on the left. Main tumor mass is marked with a dashed circle. Exemplary time-lapse imaging of a dendritic stretch over 6 days (right). White arrows point to new spines and purple arrows point to retracted spines. (R) Quantification of the dendritic turnover in connected^TUM^ neurons *in vivo* (n = 106 dendritic spines followed over time in n = 2 mice). (S) UMAP plots showing clustering of connected^TUM^ and unconnected^TUM^ neurons of sequenced co-cultures (n = 97 connected^TUM^ neurons and n = 811 unconnected^TUM^ neurons). (T) Distribution of connected^TUM^ and unconnected^TUM^ neurons across clusters showing no significant differences (n = 97 connected^TUM^ neurons and n = 811 unconnected^TUM^ neurons, Fischer test (10^5^ simulations)).

Interestingly, the application of the GABA receptor inhibitor gabazine triggered epileptiform activity of connected^TUM^ neurons in co-cultures with GB^Starter^ cells, unveiling a significant proportion of tumor cells (exceeding 96%) engaged in functional neuron-tumor networks (Figure 2E). A finding that diverges from our initial observations under physiological conditions, where neuron-glioma communication was presumed to be in the range of 10-30% of tumor cells (Figure 2G).^16^ This suggests that the manifestation of functional connectivity within these networks may require strong stimulation as in the case of neuronal hyperexcitability, occurring in later disease stages of glioblastoma,^13,14,56^ highlighting the complex relationship between structural connectivity and functional communication. We also found a strong correlation in neuronal action potential burst slopes of connected^TUM^ neurons and the GB^Starter^ response in the form of SIC half width, rise time and decay time, indicative of a sensitive functional connection (Figures 2H-J, S2A).

In summary, a majority of glioblastoma cells is functionally connected with neurons via SICs and EPSCs in the early stages of glioblastoma evolution driven by neuronal action potentials.

### Diverse neuron-tumor network formation precedes neuronal dysfunction

Next, we wanted to understand whether certain neuronal subpopulations are more prone to form neuron-tumor networks and whether this connectivity influences neuronal function and gene expression patterns. Upon examining both connected^TUM^ and unconnected^TUM^ neurons with whole-cell patch-clamp electrophysiology, we could distinguish different neuronal subtypes such as cortical regular-spiking neurons and intermittent spiking interneurons. Importantly, we found no significant differences between connected^TUM^ and unconnected^TUM^ neurons in their electrophysiological properties, including resting membrane potential, capacitance, and input resistance (Figures S2B, C). Overall, there was no difference in synaptic connectivity between connected^TUM^ and unconnected^TUM^ neurons demonstrated by the analysis of miniature excitatory (mEPSC) or inhibitory (mIPSC) postsynaptic currents (Figures 2K, S2D-F). Further, our analysis did not reveal variations in action potential firing patterns or neuronal excitability (Figures 2L, S2G). In addition, we used multielectrode array recordings and calcium imaging to demonstrate neuronal cultures with tumor cells show similar neuronal action potential burst- and firing rates and do not differ in their synchronicity to neuronal cultures with no tumor, in early stages of glioblastoma development (Figures S2H, I). Functional calcium imaging further demonstrated that both connected^TUM^ and unconnected^TUM^ neurons exhibit similar cytoplasmatic calcium transient frequencies and synchronicity (Figure 2M). Additionally, both of these neuronal populations also show co-active calcium transient patterns (Figures 2N-P, Video S2). This observation expands the concept of the neuron-tumor connectome, suggesting that connected^TUM^ neurons maintain their integration within broader neural circuits even after establishing direct connections with glioblastoma cells.

Next, we investigated whether the neuronal plasticity of connected^TUM^ neurons is affected by neuron-tumor networks. For this purpose, we employed intravital longitudinal multiphoton microscopy of patient-derived xenograft models to examine dendritic spine dynamics. Interestingly, we found dynamics comparable to physiological dendritic plasticity as previously described (Figures 2Q, R, Video S3).^57–60^

Complementing this functional investigation, we combined FACS and subsequent single-cell RNA sequencing to analyze both connected^TUM^ and unconnected^TUM^ neuronal subpopulations six days after rabies virus infection. This approach identified a consistent ratio of connected^TUM^ to unconnected^TUM^ neurons across all gene expression clusters, suggesting a widespread integration of tumor cells within neural networks irrespective of neuronal subpopulation (Figures 2S, T, S3, STAR Methods). This indicates the integration of glioblastoma cells into neural networks did not discriminate based on the functional or molecular identity of neurons, further highlighting the tumor’s ability to hijack neuronal subpopulations broadly across the brain. In summary, these data show that glioblastoma is able to connect with diverse neuronal populations preceding neuronal dysfunction.

### Neuron-tumor connectivity is patient- and cell state-dependent

As patient-specific and tumor cell-state driven heterogeneity is one hallmark of glioblastoma,^5,7,8^ we wanted to investigate tumor-intrinsic mechanisms driving neuron-tumor connectivity. For this purpose, we integrated analyses of neuron-tumor connectivity via retrograde tracing, histological tumor growth patterns of patient-derived xenografts, and single-cell RNA sequencing data from glioblastoma patients and patient-derived models. This comprehensive approach allowed us to examine the functional connectivity of these models, revealing how invasive properties of glioblastoma are associated with higher neuron-tumor connectivity.

To assess the capacity of cells from different patient-derived models to form synaptic networks, we used genes associated with the GO term for synaptogenesis^61,62^ to calculate a synaptogenic module score on single-cell RNA sequencing data (STAR Methods). In addition, we made use of a single-cell RNA sequencing-based invasivity score that was associated with invasive growth across glioblastoma patients and patient-derived models (Figures 3A, B).^8^ Interestingly, patient-derived models with a high synaptogenic score, such as S24 and T269, also show a high invasivity score (Figures 3C, D).^8^ In line with these data, patient-derived tumor models with a high synaptogenic and invasivity score also show a significantly higher mean somatokinetic speed than ones with a lower synaptogenic and invasivity score (Figures 3E, F). Our analysis indicated that tumor models with a higher propensity for invasion also exhibited greater neuronal connectivity. For this purpose, we determined the average number of connected^TUM^ neurons per GB^Starter^ cell, referred to as input-to-starter ratio. The highly invasive patient-derived models S24 and BG7 showed a mean input-to-starter ratio of approximately 40/57 +/- 8.19/12.85 respectively, whereas the less invasively growing patient-derived spheroid model P3XX showed a mean input-to-starter ratio of about 10 +/- 2.26 (Figures 3G, H, S5A, B). Furthermore, the distance distribution of connected^TUM^ neurons to GB^Starter^ cells is significantly higher in invasive patient-derived models with a mean distance of 563.8 µm +/- 2,63 in S24, 1065 +/- 7.28 µm in BG7 and 366.1 +/- 2.19 µm in P3XX (Figures 3I, J, S5C, D), suggesting a broader connectivity across larger distances. This underlines a strong correlation between synaptogenic gene expression profiles of glioblastoma across patient-derived models, their ability to integrate into neuron-tumor networks and their functionally relevant invasive cell state.

**Figure 3.**
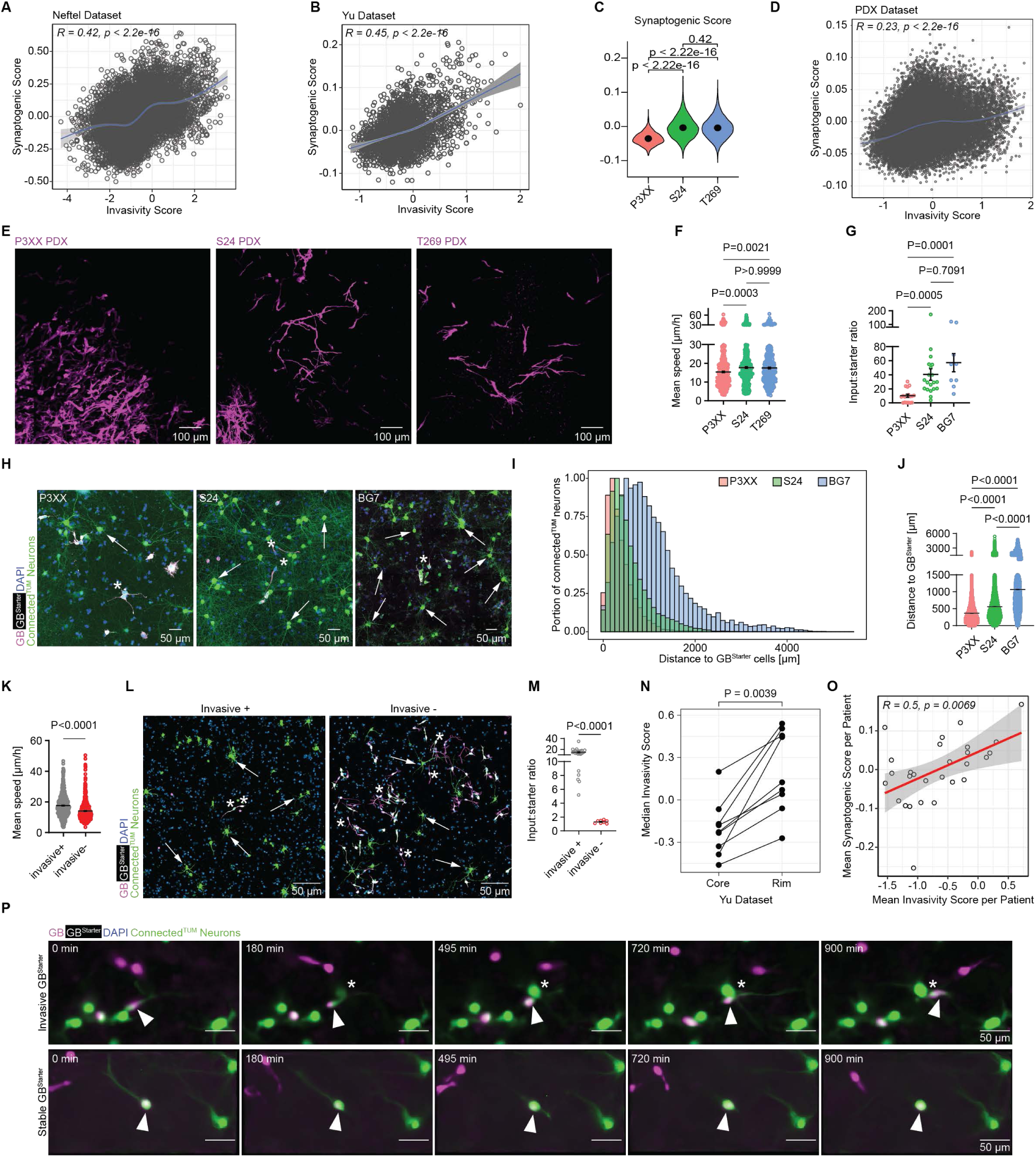
Influences of patient-and cell state-specific factors on neuron-tumor connectivity. (A) Correlation of synaptogenic and invasivity score in Neftel dataset^5^ (n = 7929 cells, Pearson’s test). (B) Correlation of synaptogenic and invasivity score in Yu dataset^63^ (n = 2795 cells, Pearson’s test). (C) Synaptogenic score compared in 3 PDX models P3XX, S24 and T269 (n = 27293 cells, Wilcoxon test). (D) Correlation of synaptogenic versus invasivity score in PDX models (n = 27293 cells, Pearson’s test). (E) *In vivo* two-photon microscopy of 3 different PDX models (P3XX (left), S24 (middle), T269 (right)) showing the invasive tumor front. Images were processed with denoise.ai. (F) Mean invasion speed of 3 different patient-derived models P3XX (left), S24 (middle) and T269 (right) in co-culture (n = 392 cells for P3XX, n = 435 cells for S24, n = 332 for T269, Kruskal-Wallis test). (G) Input-to-starter ratio comparison of 3 patient-derived models in co-culture (n = 20 (S24), n = 18 (P3XX), n = 9 (BG7) samples, Kruskal-Wallis test). (H) Representative images of retrograde tracing in patient-derived models P3XX (left), S24 (middle) and BG7 (right), showing GB^Starter^ cells (white) and their neuronal-connectome (CVS-N2c^ΔG^-eGFP(EnvA), green). Asterisks show exemplary GB^Starter^ cells and arrows point to exemplary connected^TUM^ neurons. (I) Histogram showing an overlay of the portion of connected^TUM^ neurons in relation to the distance to GB^Starter^ cells for patient-derived models P3XX, S24 and BG7 in co-culture. (J) Comparison of distance between connected^TUM^ neurons to GB^Starter^ cells in three patient-derived models in co-culture as shown in I (n = 30219 (S24), n = 17726 (P3XX), n= 10877 (BG7) cells in 3 biological replicates, one-way ANOVA). (K) Mean invasion speed shown in highly invasive microregions (DIV5-7) as compared to more stable regions (DIV12-13) in co-cultures with S24 (n = 630 cells in invasive region, n = 631 in non-invasive region, Mann-Whitney test). (L) Exemplary images of highly invasive regions (left) and less invasive regions (right). Shown are S24 GB^Starter^ cells (white, asterisks) and their connected^TUM^ neurons (CVS-N2c^ΔG^-eGFP(EnvA), green, arrows). (M) Input-to-starter ratio of highly invasive microregions (DIV5 infection) compared to less invasive, more stable microregions (DIV11 infection) in S24 co-cultures (n = 23 invasive+ and n = 8 invasive-regions, Mann-Whitney test). (N) Median invasivity score in rim versus core regions from different patients in Yu dataset^63^ (n = 2795 from 9 patients, Wilcoxon test). (O) Mean invasivity score correlated with mean synaptogenic score per patient in the Neftel dataset^5^ (n = 7929 cells from 28 patients, Pearson’s test). (P) *In vitro* live cell time-lapse imaging portraying an invasive S24 GB^Starter^ (white) and connected^TUM^ neurons (CVS-N2c^ΔG^-eGFP(EnvA), green, above). Asterix points to a newly infected, connected^TUM^ neuron adjacent to an invading GB^Starter^ cell (arrowhead). Live cell time-lapse imaging showing a stable S24 GB^Starter^ cell (white, arrowheads) and connected^TUM^ neurons (CVS-N2c^ΔG^-eGFP(EnvA), green, below). Images were processed with denoise.ai.

These results raised the intriguing question to which extent functionally distinct invasive and stationary glioblastoma cell states receive synaptic input within the same patient. For this purpose, we adopted an approach combining longitudinal, *in vivo* two-photon microscopy and serial section scanning electron microscopy to analyze synaptic connectivity across functional cell states in glioblastoma within the same patient-derived model (S24) (Figure S5E). Interestingly, this approach revealed synaptic inputs to both actively invading glioblastoma cells and to stationary tumor cells (Figures S5F, G, Video S4).^8^ To quantify whether invasive tumor cell states receive more neuronal input, we made use of combined live-cell imaging and retrograde tracing. Here, we found that more invasive tumor microregions showed a significantly higher neuron-tumor cell connectivity ratio (Figures 3K-M). These findings are in accordance with a significantly higher invasivity and synaptogenic score in the tumor rim as compared to the core within each patient, matching also the correlation of the invasivity and the synaptogenic scores per patient (Figures 3N, O, S5H).

In line with these data, we observed new infections occurring around invading GB^Starter^ cells that seem to label connected ^TUM^ neurons *en passant* via transient synaptic contacts (Figure 3P, Video S5).

These findings collectively underscored the association between a tumor cell’s synaptogenic potential at the RNA expression level with neuron-tumor connectivity and its invasiveness.

### Brain-wide recruitment of neuronal circuits by glioblastoma

Having explored the role of tumor-intrinsic factors, we examined the role of brain tumor-bearing regions on the formation of neuron-tumor networks.^25^ For this purpose, we implanted patient-derived GB^Starter^ cells into the cortex and striatum of mice, both regions frequently affected in glioblastoma patients.^64^ Investigation of patient-derived xenograft models at early stages of glioblastoma colonization (Figure S5A) revealed both long-range projections throughout the brain, including the contralateral hemisphere,^25^ as well as proximally connected^TUM^ neurons, organizing as locally connected clusters (Figures S5B, C). These long-range projections can be clearly delineated, further supported by data from co-culture models (Figure S5D). Specifically, our findings indicated that glioblastoma cells injected into the cortex exhibit more dispersed connectivity throughout the brain compared to those injected into the striatum (Figures 4A-C, S5E). While cortically localized glioblastoma showed 50% of distal connected^TUM^ neurons (defined as neuronal somata more than 1 mm away from the nearest GB^Starter^ cell), striatal tumors had 33% on average. Overall, we found approximately 9 and 14% connected^TUM^ neurons (on average 12 (+/-2)% across brain tumor regions) labeled on the contralateral hemisphere in patient-derived xenografts of cortical and striatal tumors respectively, highlighting the important role of long-range neuron-tumor networks contributing to the overall glioblastoma connectome (Figure S5F).

**Figure 4.**
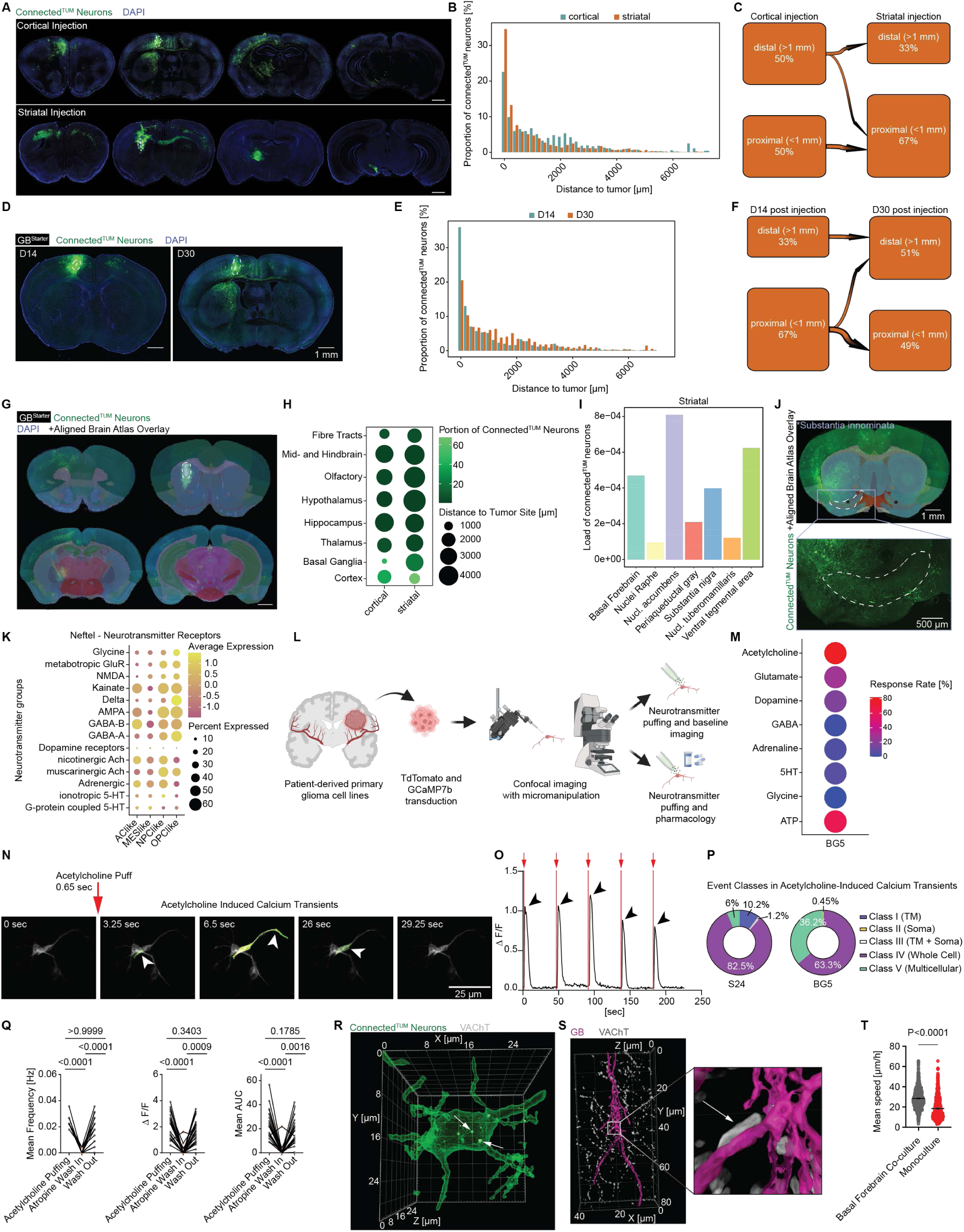
Brain tumor-bearing region-dependent formation of neuron-tumor networks. (A) Exemplary *ex vivo* brain overviews of cortical (above, SAD-B19^ΔG^-eGFP(EnvA)) and striatal (below, CVS-N2c^ΔG^-eGFP(EnvA)) tumors in PDX model S24. Dashed white lines indicate the tumor localization. Scale bar = 1 mm. (B) Histogram showing the distribution of connected^TUM^ neurons in relation to the distance from the tumor site for cortical (blue) and striatal (orange) tumors (n = 8839 connected^TUM^ neurons in n = 7 cortical tumors, n = 30528 connected^TUM^ neurons in n = 11 striatal tumors from three PDX models (S24, BG5, P3XX)). (C) River plot illustrating the distribution of distal and proximal tumor-to-neuron connections for cortical and striatal tumors (n = 8839 connected^TUM^ neurons in n = 7 cortical tumors, n = 30528 connected^TUM^ neurons in n = 11 striatal tumors from three PDX models (S24, BG5, P3XX)). (D) Representative brain sections showing the progressions of the tumor and its connectome between 14 and 30 days following tumor injection in PDX model S24 (SAD-B19^ΔG^-eGFP(EnvA)). Dashed white lines indicate the tumor localization. (E) Histogram showing the distribution of connected^TUM^ neurons in relation to the distance from the tumor site 14 (blue) and 30 (orange) days following tumor injection (n = 26419 connected^TUM^ neurons in n = 11 D14 tumors, n = 12948 connected^TUM^ neurons in n = 7 D30 tumors from three PDX models (S24, BG5, P3XX)). (F) River plot illustrating the distribution of distal and proximal neuron-tumor connections 14 and 30 days following tumor injection (n = 26419 connected^TUM^ neurons in n = 11 D14 tumors, n = 12948 connected^TUM^ neurons in n = 7 D30 tumors from three PDX models (S24, BG5, P3XX)). (G) Exemplary brain sections of PDX model S24 aligned to the Allen Brain Atlas using the QUINT workflow (STAR Methods). Connected^TUM^ neurons are shown in green (CVS-N2c^ΔG^-eGFP(EnvA)). Dashed white circle indicates the tumor localization. Scale bar = 1 mm. (H) Dot plot showing the brain region affinity of connected^TUM^ neurons depending on tumor site (n = 8839 connected^TUM^ neurons in n = 7 cortical tumors, n = 30528 connected^TUM^ neurons in n = 11 striatal tumors from three PDX models (S24, BG5, P3XX)). (I) Bar plot showing the load of connected^TUM^ neurons in various neuromodulatory circuits (n = 30528 connected^TUM^ neurons in n = 11 striatal tumors from three PDX models (S24, BG5, P3XX)). (J) Representative brain slice of PDX model S24 with connected^TUM^ neurons in the basal forebrain as shown by the alignment to the Allen Brain Atlas. The substantia innominata is marked with a dashed line. (K) Dot plot showing the expression of various neurotransmitter groups of different gene-based cell states in the Neftel dataset^5^ (n = 7929 cells). (L) Schematic workflow of the functional neurotransmitter screening in co-culture. (M) Dot plot indicating the calcium transient response rate to stimulation with different neurotransmitters (n = 78 cells from patient-derived model BG5 in n = 7 independent experiments). (N) Time-lapse imaging showing an exemplary acetylcholine puff and the following acetylcholine-induced calcium transients in a BG5 GB cell. Arrowheads point to the calcium transient. (O) Calcium imaging trace of GB cell showing acetylcholine stimulation (arrows) and the following calcium transients (arrowheads). (P) Pie charts showing the distribution of event classes in acetylcholine-induced calcium transients in two different patient-derived models (n = 166 events in n = 47 cells for S24, n = 221 events in n = 51 cells for BG5). (Q) Mean calcium event frequency, ΔF over F and area under curve (from left to right) of calcium transients in response to acetylcholine puffing, blocking through atropine and after wash-out in S24 (n = 22 cells in 2 independent experiments, Friedman test). (R) 3D rendering of a connected^TUM^ neuron (CVS-N2c^ΔG^-eGFP(EnvA), green) showing VAChT (gray, arrows) expression. (S) 3D rendering of a putative cholinergic synapse (arrow) on a GB cell (magenta) shown with staining against VAChT (gray). (T) Mean invasion speed of S24 GB cells in a basal forebrain co-culture system compared to in only S24 GB monoculture (n = 998 cells in co-culture and n = 978 cells in monoculture, Mann-Whitney test).

Our analysis also demonstrates that the proportion of distal connections of the tumor significantly increases over time, indicating a more dispersed brain-wide recruitment of neuronal circuits as the tumor progresses, diminishing the role of main tumor mass location on the spread of connected^TUM^ neurons during tumor progression (Figures 4D-F, S5G).

The quantification of connected^TUM^ neurons by brain regions showed that glioblastoma cells in both cortical and striatal regions most frequently received neuronal input from cortex, the basal ganglia and the thalamus (Figure 4H, S5H).^65–69^ While cortically localized glioblastoma cells received input mainly from isocortex, both from the ipsi- and contralateral side, striatal glioblastoma cells received most neuronal input from the basal ganglia, reflecting the high degree of neuronal connectivity within the brain regions from both cortex and basal ganglia.^65–68,70^ Interestingly, the recruited brain regions included the brainstem as pathophysiologically important regions where invasion along axonal tracts mediates lethality of glioblastoma (Figure S5H, I).^71^ Although the overall degrees of neuron-tumor connectivity is dependent on the tumor-bearing region, the overall pattern of brain-wide distribution is comparable between the cortical and striatal brain tumors. This illustrates the conserved recruitment from glioblastoma of neural circuits across brain regions (Figures 4I, S5J).

### Functional and structural acetylcholinergic neuron-tumor communication

Interestingly, distinct neuromodulatory circuits, such as those in the substantia innominata, which primarily consists of acetylcholinergic neurons,^72–74^ were recruited by glioblastoma (Figures 4I, J, S5J). To investigate the capacity of glioblastoma to directly communicate with different neuronal subpopulations, we compared co-cultures with GB^Starter^ cells and neurons from the basal forebrain, cortex and hippocampus. Interestingly, the input-to-starter ratio was not significantly different (Figures S5K, L). In line with these data, we found extensive recruitment of both glutamatergic and acetylcholinergic excitatory and GABAergic, inhibitory neurons in patient-derived xenografts and co-culture models (Figures S5M-Q). These findings highlight the tumor’s capability to integrate with various neurotransmitter systems across the brain. Supporting these findings, an unbiased analysis of publicly available single cell sequencing data^5^ showed that glioblastoma cells from human patients express genes from a broad variety of neurotransmitter receptor classes (Figures 4K, S6A). Based on the diverse neuronal subpopulations recruited by glioblastoma and neurotransmitter receptor gene expression profiles, we investigated whether different neurotransmitters released by connected^TUM^ neurons could lead to a functional response in glioblastoma cells. For this purpose, we established a functional neurotransmitter receptor screening approach (Figure 4L). Here, a targeted burst of eight neurotransmitters was sequentially applied directly onto glioblastoma cells stably expressing the genetically encoded calcium indicator GCamp7b.^75^ The resulting correlated calcium events within glioblastoma cells,^8^ triggered by a localized, time-resolved application of neurotransmitters with a high concentration similar to synaptic stimulation, served as a direct measure of functional neurotransmitter receptor expression. These calcium events could be classified into four event classes reaching from subcellular localization within a glioblastoma cell to multicellular events, as described previously.^8^ Our results indicate that glioblastoma cells are responsive to a wide range of neurotransmitters to a varying extent (Figures 4M, S6B). Interestingly, acetylcholine, ATP, glutamate and dopamine led to high degrees of responsiveness in two patient-derived models (S24, BG5), in line with the recruitment of neuromodulatory circuits and glutamatergic, excitatory neurons by glioblastoma across patient-derived xenograft models. The event areas of calcium transients after neurotransmitter response are consistently larger than those observed spontaneously, further reflected by higher rates of whole cell and multicellular transients in response to acetylcholine and glutamate (Figures 4N-P, S6C, Video S6). In contrast, GABA, serotonin and glycine showed low responsiveness of glioblastoma in both patient-derived models. The lack of functional GABA receptor expression in adult glioblastoma, as previously reported,^19^ contrasts with the structural recruitment of glioblastoma with GABAergic, inhibitory neurons. This suggests other pathways of neuron-tumor communication, potentially driving glioblastoma biology.

Next, we further characterized the functional acetylcholine receptor expression in glioblastoma cells. Here, we found that the muscarinergic acetylcholine receptor blocker atropine blocked acetylcholine-induced calcium events (Figures 4Q, S6D, E). Further investigation of single cell sequencing datasets^5,63^ revealed that the muscarinergic acetylcholine receptor M3 (CHRM3) is highly expressed in glioblastoma (Figures S6F, G).

Based on this molecular evidence, we investigated whether structural, synaptic connections between cholinergic neurons and glioblastoma cells could be detected. First, we validated that connected^TUM^ neurons indeed express the vesicular acetylcholine transporter (VAChT) and employing high-resolution light microscopy, could show there are putative cholinergic synapses directly onto the tumor cell membrane in a patient-derived xenograft model (Figures 4R, S).

Lastly, we investigated whether acetylcholinergic neurons, similar to glutamatergic neurons,^8^ could promote glioblastoma somatokinesis. Interestingly, we found that neurons from the basal forebrain promoted glioblastoma migration while monocultures of glioblastoma showed significantly slower somatokinesis, suggesting an important effect of acetylcholinergic neurons on invasive properties of glioblastoma (Figures 4T, S6H).

Taken together, local and distant neural circuits were recruited revealing extensive communication via diverse neurotransmitter receptor systems revealing acetylcholine as a functional neurotransmitter in neuron-to-glioma communication. However, the role of acetylcholine for glioblastoma biology is yet to be further characterized.

### Radiotherapy-driven remodeling of neuron-tumor networks

Increasing sequencing data of matched primary and recurrent glioblastoma samples show conflicting results regarding the role of the neural microenvironment and glioblastoma’s intrinsic neural signatures for its notorious therapeutic resistance.^11,76,77^ Exploiting time-resolved, rabies-mediated retrograde tracing, we aimed to investigate the role of neuron-tumor networks in radiotherapy-induced therapeutic resistance in a co-culture model. We found that while radiotherapeutic treatment reduced the glioblastoma cell number as expected (Figures S7A, B), the average number of connected^TUM^ neurons per glioblastoma cells significantly increased, overall increasing neuron-tumor connectivity (Figures 5A, B). We hypothesized the increased neuron-tumor connectivity is driven by neuronal activity-dependent factors and performed whole-cell patch-clamp electrophysiology of connected^TUM^ neurons with and without radiotherapy. Interestingly, we saw a significant increase in action potential bursting activity following radiotherapy, with a higher number of action potential bursts per minute and an increased area under curve of action potential bursts (Figures 5C, D). In contrast, we did not see a change in the basic electrophysiological properties of connected^TUM^ neurons after radiation, including resting membrane potential, capacitance, input resistance and rheobase (Figures S7C, D) or in their synaptic connectivity (Figures S7E-G). Taken together, the observation of increased action potential bursts after radiotherapy is in line with clinical observations of increased epileptic seizures among a subset of glioma patients following radiotherapy.^78^

**Figure 5.**
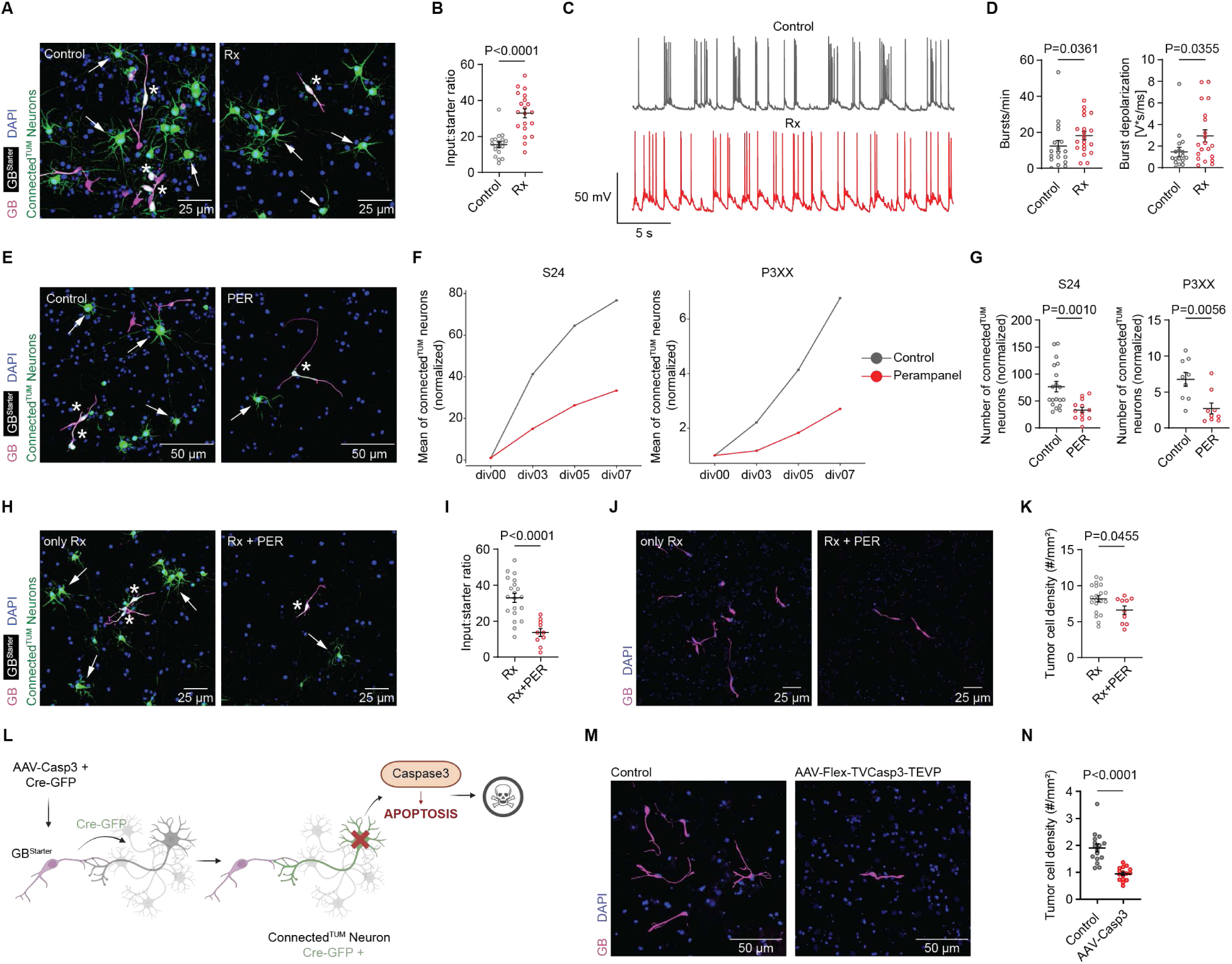
Tackling neuron-tumor networks with rabies and combined radiotherapy and neuronal activity inhibition. (A) Probability maps of S24 GB^Starter^ cells (white, asterisks) and connected^TUM^ neurons (CVS-N2c^ΔG^- eGFP(EnvA), green, arrows) in co-culture under control conditions (left) versus after radiation therapy with 4 Gray (right). (B) Input-to-starter ratio under control compared to radiotherapy conditions (n = 19 control versus 20 radiotherapy-treated samples, Mann-Whitney test). (C) Representative whole-cell current-clamp recordings of spontaneous burst firing in connected^TUM^ neurons under control conditions (top) versus after radiotherapy (bottom). (D) Bursts per minute (left panel, n = 18 control and n = 20 irradiated neurons, Mann-Whitney test) and normalized burst area (right) panel, n = 18 control and n = 20 irradiated neurons, Mann-Whitney test). (E) Exemplary probability maps of the neuronal connectome of glioblastoma under control condition (left) versus perampanel treatment (right) showing GB^Starter^ cells (white, asterisks) and connected^TUM^ neurons (CVS-N2c^ΔG^-eGFP(EnvA), green, arrows). (F) Normalized mean count of connected^TUM^ neurons in S24 (left) and P3XX (right) GB patient-derived models over a time course of 7 days under control conditions and perampanel treatment (n = 19 control versus 13 perampanel-treated samples in S24, n = 9 control versus n = 9 perampanel-treated samples in P3XX). (G) Quantitative comparison of normalized mean count of connected^TUM^ neurons in S24 (left) and P3XX (right) on day 07 of treatment (n = 19 control versus 13 perampanel-treated samples in S24, n = 9 control versus n = 9 perampanel-treated samples in P3XX, Mann-Whitney test). (H) Probability maps of GB^Starter^ cells (white, asterisks) and connected^TUM^ neurons (CVS-N2c^ΔG^- eGFP(EnvA), green, arrows) in only irradiated (left) versus simultaneous irradiation and perampanel-treatment (right) conditions. (I) Input-to-starter ratio in only irradiated versus simultaneous irradiation and perampanel-treatment conditions (n = 20 only irradiated versus 10 irradiated and perampanel-treated samples, unpaired t-test). (J) Representative images of tumor regions in only irradiated (left) compared to simultaneous irradiation and perampanel treatment (right) conditions in S24 co-culture. (K) Tumor cell density in cell count per mm^2^ under only irradiated conditions compared to a combination of irradiation and perampanel-treatment in co-culture (n = 20 only irradiated versus 10 irradiated and perampanel-treated samples, unpaired t-test). (L) Schematic of experimental paradigm for genetic ablation of connected^TUM^ neurons. (M) Representative images of *in vitro* S24 GB cells under control conditions (left) compared to the genetic ablation of connected^TUM^ neurons (right). (N) Quantification of S24 tumor cell density in cell count per mm^2^ under control conditions compared to the genetic ablation of connected^TUM^ neurons in co-culture (n = 16 control compared to n = 16 caspase treated samples, Mann-Whitney test).

Next, we investigated whether neuron-tumor connectivity is driven by neuronal activity, similar to synaptogenesis in neuron-to-neuron synapses.^79,80^ For this purpose, we employed the non-competitive AMPAR antagonist perampanel (PER), commonly used as antiepileptic drug to inhibit neuronal activity.^81^ In consequence, the neuron-tumor connectivity and tumor cell number significantly decreased, highlighting the role of intrinsic neuronal activity in the formation of neuron-tumor networks (Figures 5E-G, S7H).^8,16,17^

These data led us to the question whether simultaneous inhibition of neuronal activity and radiotherapy would decrease neuron-tumor network connectivity and increase therapeutic efficacy.

Interestingly, we saw that neuron-tumor connectivity is significantly reduced after combined radiotherapy and AMPAR inhibition as compared to radiotherapy alone (Figures 5H, I). In consequence, we could also see how glioblastoma progression was reduced by this therapy combination, indicative of neuron-to-glioma synaptic communication contributing to therapeutic resistance (Figures 5J, K).

Taken together, radiotherapy-induced neuronal activity promotes neuron-tumor connectivity with combined inhibition of neuronal activity and radiotherapy showing increased therapeutic effects, requiring further clinical-translational investigation.

### Rabies virus-based ablation of tumor-connected neurons inhibits glioblastoma progression

In addition to pharmacological perturbation of neuron-tumor networks, we investigated whether retrograde tracing with the modified rabies virus itself could be used to specifically ablate both local and distant connected^TUM^ neurons to inhibit glioblastoma progression. For this purpose, we implemented a Cre-loxP strategy to specifically ablate connected^TUM^ neurons in a co-culture model. Thus, we infected the neural tumor microenvironment with an AAV expressing a Cre-dependent genetically engineered designer caspase 3, a caspase whose activation drives cells to apoptosis. Hereby, we could specifically kill connected^TUM^ neurons and investigate its effect on tumor cells (Figure 5L).^82,83^ The eradication of connected^TUM^ neurons across all neuronal subtypes resulted in a significant reduction of tumor cells as compared to controls (Figures 5M, N).

Taken together, these data demonstrate how a modified rabies virus could in principle be used to kill heterogeneous connected^TUM^ neuronal subpopulations via retrograde infection and thus inhibit glioblastoma growth.

## DISCUSSION

It is becoming increasingly clear that synaptic neuron-tumor networks are an important hallmark of yet incurable glioblastomas.^12,13^ Our research introduces a comprehensive and reproducible methodology platform capable of investigating the neuronal connectome of glioblastoma across a range of models, from purely human tissue models over co-culture models of neurons and tumors to patient-derived xenografts. Technologies such as monosynaptic retrograde tracing are especially important in the context of highly invasive tumors such as glioblastoma. This contrasts sharply with traditional dye injection techniques that could get taken up by neurons nearby tumor cells, precluding cellular specificity of the labeled neuronal connectome.^29^ Importantly, the ability to investigate the neuronal connectome of patient-derived models in a human tissue context opens up the potential for personalized therapeutic approaches, disconnecting neuron-tumor networks.

By integrating longitudinal imaging, electrophysiology, molecular characterization, and functional tumor biological assays, we have gained important insights into the malignant circuitry’s evolution. A nuanced picture has emerged, revealing the bidirectional mechanisms that underpin neuron-tumor connectivity: tumor cells establish transient, functional connections with neurons regardless of their molecular or functional properties. Concurrently, the functional connectivity between neurons and tumor cells can be significantly increased by neuronal activity.

The impact of known glioma-induced alterations in neural circuits^17,24,46,49,84^ on overall brain function and their contribution to disease advancement warranted further investigation. Our findings showed that, at least in the initial stages of glioblastoma colonization, the functional and molecular properties of neurons remain unchanged, potentially setting the stage for neuronal hyperexcitability as the disease progresses. Thus, these findings together suggest a model where the establishing neuron-tumor networks precedes neuronal dysfunction in the course of the disease. This would also be in concordance with clinical findings where epileptic seizures occur in later disease stages where curable surgical resection of glioblastoma is no longer feasible.^56^ Functional imaging of connected^TUM^ and unconnected^TUM^ neurons revealed that connected^TUM^ neurons are well integrated into neural circuits of unconnected^TUM^ neurons, as evidenced by co-active firing patterns. With neuronal activity being able to elicit calcium transients in glioblastoma cells, this suggests the concept of a primary, directly connected and secondary, indirectly connected neuronal connectome. These data also make it unlikely that connected^TUM^ neurons are created via neurogenesis, a phenomenon that has been previously described in prostate cancer,^85^ as neurons derived from neurogenesis presumably need several weeks of integrating into neuronal networks.^86^ These complex networks highlight the importance of investigating bidirectional interactions between glioblastoma and the central nervous system across scales, including distant and even non-tumor connected brain regions. The specificity of neural influence, especially how particular neuronal types and neurotransmitters distinctively affect various cancer types, remains an area for further investigation.

Our work highlights the readiness of tumor cells to engage in functional communication with various neuronal subpopulations across the brain including various neuromodulatory circuits by expressing a spectrum of neurotransmitter receptors. Specifically, we found functional, muscarinergic acetylcholine receptors on glioblastoma cells across patient-derived models and structural acetylcholinergic neuron-to-glioma synaptic contacts. Further, we found that acetylcholinergic neurons promote glioblastoma progression.

Interestingly, not all neurotransmitter receptors were functionally expressed on glioblastoma cells, implying that there might be either structurally present and functionally “silent” neuron-glioma synapses. In the case of structurally connected GABAergic neurons, co-transmitted neurotransmitters such as acetylcholine might play an important role,^87^ or paracrine neuron-tumor communication^88,89^ could mediate tumor biological effects.

Importantly, early synaptic connections to brainstem neurons hinted at a strategy for glioblastoma invasion along axonal pathways into the brainstem, a critical factor in the disease’s lethality.^71^ This observation suggests that potentially initial synaptic connections prompted glioblastoma’s migration along axonal structures, paving a path for invasion into distant brain regions. The importance of neuron-glioma synapses for distant invasion is further supported by an increased neuron-tumor connectivity in more invasively growing tumors and invasive tumor cell states, driven by their synaptogenic gene expression profiles.

In addition to the molecular and functional characterization of neuron-tumor networks, retrograde tracing in the context of glioblastoma enabled the investigation of how neuron-tumor networks are formed and therapeutically exploited. Interestingly, we could see how neuronal activity-dependent formation of neuron-tumor networks parallels similar establishments of physiological synaptic connections during development.^79,80^ Furthermore, we could see that increasing neuronal activity through radiotherapy increased neuron-tumor connectivity and show an inhibition of AMPA receptors in combination with standard-of-care radiotherapy yields synergistic therapeutic effects. This demonstrates an additional role of neuron-glioma synaptic communication in therapeutic resistance, explaining a potential role of neuronal gene expression signatures of glioblastoma in the recurrent setting.^4^

Using our rabies-based tracing approach, we demonstrate how this system could in principle be directly used to induce apoptosis specifically in neurons connected to tumor cells, thereby decreasing tumor progression. Our proof-of-concept investigation in several model systems enables further modification of rabies virus constructs to not only eliminate cancer cells but also their associated neuronal connectome as a potential novel therapeutic strategy. Such a viral approach to target glioblastoma could even be adapted to specifically disconnect neuron-tumor network connectivity, adding to other promising immunotherapeutic, viral strategies tackling not only glioblastoma directly but its associated neuronal connectome.^90^

Taken together, we established a novel framework to investigate the neuronal connectome of glioblastoma that can be translated to study not only other brain tumors but also cancers outside the brain. Using this scalable technology, we furthered our understanding about the organization, formation and therapeutic opportunities yielded by neuron-tumor networks enabling further investigation.

### Limitations of the study

This study introduces a technology platform allowing a comprehensive multimodal look into the neuron-tumor connectome in glioblastoma, employing a rabies virus-based retrograde tracing system to explore these complex interactions. Despite the insights provided on a basic science and clinical-translational level, there are limitations that merit consideration for a more comprehensive understanding and broader application of the findings. The study primarily focuses on the early stages of neuron-tumor network formation, highlighting a need for further exploration across various stages of tumor development to fully understand how these interactions evolve and impact disease progression and therapeutic responses over time. One limitation of the experimental platform is the neurotoxic potential associated with the use of rabies virus for retrograde tracing over time.^91–93^ This issue underscores the importance of utilizing and further adapting less toxic rabies-based labeling strategies for glioblastoma,^93,94^ to enable longer observation periods without adverse effects on neuronal health. Further, rabies virus-mediated retrograde tracing did not label all synaptic inputs in previous work, illustrating that the labeled connected^TUM^ are possibly still an underestimation of the entire neuronal connectome of glioblastoma.^95,96^ Additionally, while a high level of neuron-tumor connectivity is observed, the precise mechanisms underpinning the synaptic interactions between neurons and glioma cells, especially regarding the role of neuronal action potential-driven slow inward currents (SICs), remain unclear and require further elucidation. While we demonstrate a biological effect of acetylcholinergic neurons on glioblastoma biology, further investigation of acetylcholinergic neurotransmission as well as the specific effects of other neuronal subpopulations on glioblastoma is needed.

Furthermore, the therapeutic effects observed between radiotherapy and neuronal activity inhibition via perampanel present a promising therapeutic avenue, warranting validation across diverse model systems and in clinical trials to confirm their potential. The feasibility of using a modified rabies virus to specifically ablate connected^TUM^ neurons also poses a significant opportunity. While the study provides a proof-of-concept, further research is necessary to determine how these viral constructs can be adapted for efficacy and safety in a clinical-translational context without the need for genetically modifying neurons via AAVs. Lastly, the application of retrograde tracing in patient-derived glioblastoma spheroids suggests the potential for this methodology to be extended to other types of cancer, both within and outside the brain. Investigating whether other tumors receive synaptic input and how neuron-tumor interactions vary across different malignancies could open new paths for cancer research, enhancing our understanding of these complex networks and paving the way for novel therapeutic strategies across oncology.

**Supplementary Video 1:** Dynamic investigation of single tumor cell and whole tumor-associated neuronal connectome in glioblastoma, related to Figure 1. Shown are sparse and dense labeling approaches of glioblastoma in co-culture of neurons and tumor cells, depicting different modalities of rabies-based retrograde tracing.

**Supplementary Video 2:** Functional networks of connected^TUM^ and unconnected^TUM^ neurons during early glioblastoma colonization shown with simultaneous calcium imaging in co-cultures, related to Figure 2. Connected^TUM^ neurons are embedded in a synchronously firing network of unconnected^TUM^ neurons.

**Supplementary Video 3:** Structural plasticity of dendritic spines in connected^TUM^ neurons shown with *in vivo* two-photon microscopy of glioblastoma, related to Figure 2. High-resolution time-lapse imaging of connected^TUM^ neurons showing physiological dendritic plasticity.

**Supplementary Video 4:** Neuron-to-glioma synapse reconstructions across functional cell states in a patient-derived xenograft model (S24), related to Figure 3. Correlation of light and electron microscopy reveals synaptic input on invasive and stable glioblastoma cells.

**Supplementary Video 5:** Investigating the dynamic neuron-tumor connectome with longitudinal imaging, related to Figure 3. *In vitro* live cell time-lapse imaging portraying an *en passant* infection of a connected^TUM^ neuron by an invasive GB^Starter^ cell in contrast to a stable GB^Starter^ cell within the same time frame.

**Supplementary Video 6:** Functional muscarinergic acetylcholinergic receptor expression of patient-derived glioblastoma cells, related to Figure 4. Shown are glioblastoma cells responding to acetylcholine stimulation in the form of subsequent calcium transients in co-culture of glioblastoma cells and neurons.

**Supplementary Table 1:** Overview of patient-derived glioblastoma models

**Supplementary Table 2:** Overview of genes included in neurotransmitter group analysis of single-cell RNA sequencing data

## METHODS

### KEY RESOURCES TABLE

| REAGENT or RESOURCE | SOURCE | IDENTIFIER |
| --- | --- | --- |
| Antibodies |  |  |
| Mouse monoclonal anti-beta-III-tubulin | Abcam | Cat#ab7751,<br>RRID:AB_306045 |
| Guinea pig polyclonal anti-NeuN | Synaptic Systems | Cat#266004,<br>RRID:AB_2619988 |
| Rabbit polyclonal anti-VACHT | Synaptic Systems | Cat#139103,<br>RRID:AB_887864 |
| Chicken polyclonal anti-S100B | Synaptic Systems | Cat#287006,<br>RRID:AB_2713986 |
| Rat monoclonal anti-MBP | Novus Biologicals | Cat#NB600-717,<br>RRID:AB_2139899 |
| Rabbit polyclonal anti-Iba1 | FUJIFILM Wako<br>Pure Chemical<br>Corporation | Cat# 019-19741,<br>RRID:AB_839504 |
| Mouse monoclonal anti-Nestin | Abcam | Cat#ab22035<br>RRID:AB_446723 |
| Chicken polyclonal anti-GFP | Abcam | Cat#ab13970<br>RRID:AB_300798 |
| Chicken polyclonal anti-GFP | Aves Labs | Cat#GFP-1020,<br>RRID:AB_1000024<br>0 |
| Rabbit polyclonal anti-RFP, pre-adsorbed | Rockland | Cat#600-401-379,<br>RRID:AB_1118280<br>7 |
| Rabbit polyclonal anti-mCherry | Abcam | Cat#ab167453,<br>RRID:AB_2571870 |
| Guinea pig polyclonal anti-RFP | Synaptic Systems | Cat#390004,<br>RRID:AB_2737052 |
| Mouse monoclonal anti-CAMK2 | Abcam | Cat#ab22609,<br>RRID:AB_447192 |
| Rabbit polyclonal anti-ChAT | invitrogen | Cat#PA5-29653,<br>RRID:AB_2547128 |
| Rabbit polyclonal anti-Parvalbumin | Abcam | Cat#ab11427,<br>RRID:AB_298032 |
| Goat anti-chicken Alexa488 | invitrogen | Cat#1458638<br>and 2304258,<br>RRID:AB_2534096 |
| Goat anti-rabbit Alexa647 | invitrogen | Cat#1981173 and<br>2299231,<br>RRID:AB_2535813 |
| Goat anti-guinea pig Alexa647 | invitrogen | Cat#A-21450,<br>RRID:AB_141882 |
| Goat anti-guinea pig Alexa546 | invitrogen | Cat#A11074,<br>RRID:AB_2534118 |
| Goat anti-chicken Alexa647 | invitrogen | Cat#A-21449,<br>RRID:AB_2535866 |
| Goat anti-rat Alexa647 | invitrogen | Cat#A-21247,<br>RRID:AB_141778 |
| Goat anti-rabbit Alexa568 | invitrogen | Cat#A11011,<br>RRID:AB_143157 |
| Goat anti-mouse Alexa647 | invitrogen | Cat#A21235,<br>RRID:AB_2535804 |
| Goat anti-mouse Alexa568 | invitrogen | Cat#A11011,<br>RRID:AB_144696 |
| Goat anti-chicken Alexa488 | invitrogen | Cat#A32931,<br>RRID:AB_2762843 |
| Streptavidin Alexa647 conjugate | Thermo Fischer | Cat#S21374,<br>RRID:AB_2336066 |
| **Bacterial and virus strains** |  |  |
| RABV CVS-N2C(deltaG)-EGFP | addgene | #73461 |
| SAD-B19(deltaG)-EGFP | addgene | #32634 |
| pAAV-mDlx-NLS-mRuby2 | addgene | #99130 |
| pAAV-flex-taCasp3-TEVp | addgene | #45580 |
| pSADdG/CreGFP | Charité Viral Vector<br>Core | BR-26 |
| pAAV.Syn.NES-jRGECO1a.WPRE.SV40 | addgene | #100854 |
| **Biological samples** |  |  |
| Patient-derived xenografts (PDX) | This paper | N/A |
| **Chemicals, peptides, and recombinant proteins** |  |  |
| Poly-L-lysine | Sigma | P4707 |
| Neurobasal Medium | Gibco | 11570556 |
| Hibernate™-A Medium | Gibco | A1247501 |
| B-27 Supplement for neuronal co-culture | Gibco | 17504044 |
| B27 supplement without vitamin A for GB culture | Gibco | 12587010 |
| L-glutamine (GlutaMAX™-I (100x)) | Gibco | 35050038 |
| Antibiotic-Antimycotic (100x) | Gibco | 15240062 |
| MgSO<sub>4</sub> | Sigma Aldrich | M3409 |
| DMEM/F12 Medium | Gibco | 11330032 |
| Insulin solution human | Sigma | I9278 |
| Heparin | Sigma | H4784 |
| EGF | Biotechne | 236-EG-200 |
| FGF Recombinant Protein | Life Technologies | PHG0021 |
| FGF Recombinant Protein 1mg | Life Technologies | PHG0023 |
| Accutase Solution | FisherScientific | A1110501 |
| 0.05% Trypsin-EDTA (1x) | Gibco | 25300-054 |
| 2.5% Trypsin (10x) | Gibco | 15090-046 |
| NGF | Miltenyi Biotech | 130-127-430 |
| mTeSR™ Plus | StemCell<br>Technologies | #100-0276 |
| ReLeaSR | StemCell Technologies | #05872 |
| 1% N2 supplement | Gibco | 17502048 |
| Minimum Essential Medium from non-essential amino acids | Gibco | 11140050 |
| Laminin | ThermoFischer | 23017015 |
| BDNF | Peprotech | #450-02 |
| NT-3 | Peprotech | #450-03 |
| Doxycycline-HCl | FisherScientific | 15473189 |
| Cytosine arabinoside | Sigma | #C6645 |
| HyClone FBS | Cytiva | SH30071.03HI |
| FBS | Anprotec | AC-SM-0041 |
| AcX | invitrogen | A20770 |
| Ammoniumpersulfate | Sigma | A3678 |
| TEMED | Merck | T9281 |
| 4-Hydroxy-TEMPO | Merck | 176141 |
| Acrylamide | Sigma | A9099 |
| Sodiumacrylate | Sigma | 408220 |
| Neurobiotin Tracer | Vector Laboratories | SP-1120 |
| HEPES | Sigma Aldrich | 7365-45-9 |
| HEPES solution | Sigma Aldrich | H0887 |
| D-Glucose | Sigma Aldrich | G8644 |
| N-Methyl-D-Glucamin | Sigma Aldrich | M2004 |
| ASC acid | Riedel-de-Haën | 33034 |
| HI horse serum | ThermoFischer | 26050-070 |
| N,N'-Methylenbisacrylamide | Merck | 146072 |
| Perampanel | BioCrick | BCC1847;<br>CAS: 380917-97-5 |
| Glutamate | Sigma Aldrich | G8415 |
| Acetylcholine | Sigma Aldrich | A6625 |
| 5-HT | Sigma Aldrich | 14927 |
| GABA | Sigma Aldrich | A2129 |
| Adrenaline | Tocris | 5169 |
| Dopamine hydrochloride | Sigma Aldrich | H8502 |
| Glycine | Sigma Aldrich | G7126 |
| Alexa Fluor 594 hydrazide sodium salt | invitrogen | A10442 |
| Atropine sodium salt | Sigma Aldrich | A0132 |
| Dulbecco's Phosphate Buffered Saline | Sigma Aldrich | D8537 |
| Millicell cell culture inserts 0.4 um | Merck | PICM0RG50 |
| CellTiter-Glo® Luminescent Cell Viability Assay | Promega | G7570 |
| NaCl | Sigma-Aldrich | S7653 |
| KCL | Fluka | 60129 |
| NaHCO3 | Fluka | 71627 |
| NaH2PO4 | Fluka | 71496 |
| Triton™ X-100 | Sigma-Aldrich | T9284 |
| L-glutamine (GlutaMAX™-I (100x)) | Gibco | 35050-038 |
| Evans Blue | Sigma | E2129 |
| SlowFade™ Gold Antifade Mountant | ThermoFischer | S36936 |
| DAPI | Sigma | D9542 |
| PFA 4% | Roth | P087.3 |
| Sulforhodamine 101 (SR101) | Sigma Aldrich | S7635 |
| Tetramethylrhodamine isothiocyanate dextran (TRITC) | Sigma Aldrich | 52194 |
| Cacodylic acid-Na-salt-3H2O | Serva | 15540.1 |
| Potassium ferricyanide | Serva/Sigma | 107H3450 |
| Osmium tetroxide | Serva | 31253.04 |
| Glycid Ether 100 for electron microscopy | Serva | 21045.02 |
| MNA hardener | Carl Roth | 8639.2 |
| DMP-30 | Carl Roth | 8621.1 |
| DBA Härter | Carl Roth | 8623.2 |
| DDSA | Serva | 20755.02 |
| Propylenoxide | VWR Chemicals | 27165.295 |
| Uranyl acetate | Serva | 77870 |
| SEM pin stub (0,5"/6mm length) | Agar scientific | #G301F |
| Acheson silver | Plano | #3692 |
| Experimental models: Cell lines |  |  |
| S24 | This paper | N/A |
| BG5 | This paper | N/A |
| BG7 | This paper | N/A |
| P3XX | This paper | N/A |
| T269 | This paper | N/A |
| GG16 | This paper | N/A |
| U3017MG | Xie et al., 2015 | N/A |
| U3085MG | Xie et al., 2015 | N/A |
| U3047MG | Xie et al., 2015 | N/A |
| U3048MG | Xie et al., 2015 | N/A |
| Experimental models: Organisms/strains |  |  |
| NMRI-Foxn1 nu/nu | Charles River and Janvier | BL210203171 |
| WISTAR | Janvier | N/A |
| C57BL/6 | Janvier | N/A |
| WA01/H1 | WiCell | N/A |
| HD6 | Heidelberg University, Heidelberg, Germany | N/A |
| Recombinant DNA |  |  |
| Plasmid: pFU-TVA-2A-mCherry-2A-oGlycoprotein | addgene | #85225 |
| Plasmid: mGFP | De Paola et al., 2003 | N/A |
| Plasmid: GCamp7-tdTomato | Dana et al., 2019 | N/A |
| Plasmid: GFP | Osswald et al., 2015 | N/A |
| Plasmid: tdTomato | Osswald et al., 2015 | N/A |
| Software and algorithms |  |  |
| Fiji | Schindelin et al., 2012 | https://imagej.nih.gov/ij/ |
| NIS-Elements AR Analysis 5.41.00 64-bit | Nikon | N/A |
| Arivis Vision4D 3.5.0 | arivis AG, Munich, Germany | https://imaging.arivis.com/en/imaging-science/arivis-vision4d |
| R Studio 1.4 | RStudio Team, 2023 | N/A |
| GraphPad Prism Version 9 | GraphPad | N/A |
| Adobe Illustrator 28.2 64-bit | Adobe | N/A |
| Ilastik 1.4.0 | Berg et al., 2019 | https://www.ilastik.org/development.html |
| AquA | Wang et al., 2019 | N/A |
| PATCHMASTER Igor Pro 6.21 | HEKA | N/A |
| DaVinciResolve 18 | Blackmagicdesign | https://www.blackmagicdesign.com/de/products/davinciresolve/ |
| TrakEM | Cardona et al., 2012 | N/A |
| Zen Blue 3.5 | Zeiss |  |
| Zen Black 2.3 | Zeiss |  |
| Leica Application Suite X | Leica Microsystems CMS GmbH | https://www.leica-microsystems.com/de/produkte/mikroskop-software/p/leica-las-x-ls/ |
| MATLAB 9.120.1884302 | MathWorks |  |
| Easy Electrophysiology |  | RRID:SCR_021190 |
| pClamp 11 | Molecular Devices |  |
| Autodesk 3ds max | Autodesk | https://www.autodesk.de/products/3ds-max/overview |
| Single-cell RNA-sequencing datasets |  |  |
| Human single-cell RNA-sequencing glioblastoma dataset | Neftel et al., 2019 | GSE131928 |
| Human single-cell RNA-sequencing glioblastoma dataset | Yu et al., 2020 | GSE117891 |
| Xenograft single-cell RNA-sequencing glioblastoma datasets | Hai et al., 2021 | N/A |
| Human single-cell RNA-sequencing glioblastoma dataset | Ruiz-Moreno et al., 2022 | GSE141946, GSE166418, GSE162631, GSE154795, GSE141383, GSE182109, GSE173278 |
| Other |  |  |
| Multirad 225 X Ray Irradiation System | Faxitron | BLE1900269 |
| FACSAria Fusion 2 Bernhard Shoor | BD | N/A |
| FACSAria Fusion Richard Sweet | BD | N/A |

#### Human specimens and animal models

Human tissues used for organotypic slice cultures were obtained after approval of the local regulatory authorities (ethical codes 23-1233-S1, 23-1234-S1 and S-005/2003). Human patient samples were pseudonymized manually.

Male NMRI nude mice were used for all animal studies involving patient-derived glioblastoma models. All animal procedures were performed in accordance with the institutional laboratory animal research guidelines following approval of the Regierungspräsidum Karlsruhe, Germany. Efforts were made to minimize animal suffering and reduce the number of animals used according to the 3R principles. Mice were clinically scored and if they showed marked neurological symptoms or weight loss exceeding 20%, experiments were terminated. No maximum tumor size was defined for the invasive brain tumor models.

#### Lentiviral vector and plasmid generation of pFU-TVA-2A-mCherry-2A-oGlycoprotein

To generate lentiviruses expressing, EnvA TVA receptor (TVA), rabies glycoprotein (oG), and mCherry, we sub-cloned TVA-2A-mCherry-2A-oGlycoprotein into a lentiviral vector (‘pFU-‘) using In-Fusion cloning (Takara). TVA-2A-mCherry-2A-oGlycoprotein was amplified from p306 (Zurich virus core), and cloned into a pFU vector using ECORI and BAMHI sites.

#### Packaging of CVS-N2c^ΔG^ and SAD-B19^ΔG^

Rabies viruses used in this study were produced as described previously.^97^ Briefly, B7GG cells were transfected by Lipofectamine 3000 (Thermo Fischer) with rabies virus genomic vectors RabV CVS-N2c^ΔG^-eGFP (Addgene plasmid #73461) or SAD-B19^ΔG^-eGFP (modified from Addgene plasmid # 32634). Supernatant was collected over several days and the recovered virus was re-transfected in B7GG cells for a final collection step. For pseudotyping, the supernatant containing unpseudotyped viruses and the rabies with the envelope protein EnVA of the Avian Sarcoma and Leukosis virus were applied on BHK-EnVA cells. 3-5 days later, the EnVA-pseudotyped rabies virus was collected, filtered and concentrated using an ultracentrifuge. Titer was determined by infection of HEK293T-TVA cells with serially diluted viruses. RabV CVS-N2c(deltaG)-EGFP was a gift from Thomas Jessell (Addgene plasmid #73461; http://n2t.net/addgene:73461; RRID: Addgene_73461). pSADdeltaG-F3 was a gift from Edward Callaway (Addgene plasmid #32634; http://n2t.net/addgene:32634; RRID: Addgene_32634).

#### Patient-derived glioblastoma cultures

Patient-derived glioblastoma spheroid models from resected tumor material were cultivated as previously described^8,9,98^ in DMEM/F-12 under serum-free, non-adherent, ‘stem-like’ conditions, which includes B27 supplement without Vitamin A, insulin, heparin, epidermal growth factor, and fibroblast growth factor as described before.^99^ Glioblastoma models U3085MG, U3048MG, U3047MG, U3017MG were obtained from the Human Glioma Cell Culture (HGCC, www.hgcc.se) biobank resource at Uppsala University, Uppsala, Sweden.^51^

The patient-derived glioblastoma spheroid models were transduced with lentiviral vectors for the TVA receptor with a modified TVA-P2A-mCherry-2A-oG construct based on the Addgene plasmid #85225, membrane-bound GFP with the pLego-T2-mGFP construct,^100^ and for calcium imaging with the pLego-T2-GCaMP7b-tdTomato construct.^75^ For direct labeling, TVA-oG-mCherry expressing glioblastoma spheroids were transduced with rabies virus constructs SAD-B19^ΔG^- eGFP(EnVA) or CVS-N2c^ΔG^-eGFP(EnVA) (Addgene #73461) prior to further experiments.^33^ Transduced cells were sorted regularly by FACS with either FACSAria Fusion 2 Bernhard Shoor or FACSAria Fusion 1 Richard Sweet. Following filters were used for the respective fluorophores: 610/20 for mCherry, 530/30 for GFP, 586/15 for tdTomato.

#### 850k methylation array analysis

The Illumina Infinium Methylation EPIC kit was used to obtain the DNA methylation status at >850,000 CpG sites in patient-derived glioblastoma spheroid models, according to the manufacturer’s instructions at the Genomics and Proteomics Core Facility of the German Cancer Research Center in Heidelberg, Germany, as described previously.^101^ The molecular classification of patient-derived glioblastoma models used in this study can be found in Table S1.

#### Harvesting cortical tissue from human patients

During surgical interventions, cortical tissue proximal to deeper pathologies was precisely and safely extracted, guided by neuro-navigation techniques. To ensure the removal of non-damaged tissue, we applied a refined method recently detailed by our group.^102^ This technique enhances the accurate identification and collection of cortical tissues, aiming to minimize harm. The criteria for selecting human slice cultures are rigorously defined to maintain the material’s study relevance and integrity. Specifically, tissue designated for slice culture is required to be more than 10 millimeters away from identified pathologies, like metastases or vascular issues, establishing a safety margin to exclude potentially compromised tissue not evident visually. For glioma tumors, the criteria are stricter, demanding over 20 millimeters of separation from the tumor, acknowledging gliomas’ diffuse infiltration potential. While ensuring 100% pathology-free tissue is challenging, we leveraged Scattered Raman Histology and AI-based detection to mitigate the impact of any significant tumors or pathologies on the harvested cortex in selected patients.^103^

#### Human organotypic slice cultures

Human neocortical slices were prepared following a recently described procedure.^104–107^ Immediately after resection, cortical tissue was transported to the laboratory in a carbogen-saturated “Preparation medium” (Gibco Hibernate^TM^ media with 0.5 mM Gibco GlutaMax^TM^, 13 mM Glucose, 30 mM NMDG, 1% Anti-Anti, 1 mM ASC Acid, and HI Horse Serum) on ice. Under a 10x microscope, capillaries and damaged tissue were microdissected, and the arachnoidia was microsurgically removed. The collection medium, enriched with GlutaMax and NMDG, ensured optimal tissue recovery. Cortical slices, 300 μm thick, were created using a vibratome (VT1200, Leica Germany) and incubated in the preparation medium for 10 minutes pre-plating to minimize variability from tissue trauma. Typically, tissue blocks (1 cm × 2 cm) allowed for 15 sections, with 1-3 sections per insert being carefully spaced. A polished wide-mouth glass pipette facilitated slice transfer. The slices were then maintained in a growth medium composed of Neurobasal L-Glutamine (Gibco) supplemented with 2% serum-free B-27 (Gibco), 2% Anti-Anti (Gibco), 13 mM d-glucose (Sigma-Aldrich), 1 mM MgSO4 (Sigma-Aldrich), 15 mM Hepes (Sigma-Aldrich), and 2 mM GlutaMAX (Gibco). The medium was refreshed 24 hours after plating and then every 48 hours. For inoculation, target cells were prepared as previously mentioned, undergoing post-trypsinization centrifugation, harvesting, and resuspension in PBS at 20.000 cells/μl. Cells were inoculated into tissue sections using a 10 μL Hamilton syringe to deliver 1 μL onto the white matter, then incubated at 37°C for a week with medium changes every 48 hours. Tumor proliferation was assessed using fluorescence imaging with an inverted microscope (Observer D.1; Zeiss). After the designated culture period, sections were fixed for immunostaining.

#### Surgical procedures

For *in vivo* two-photon imaging, surgical procedures were performed as described previously.^8,9,16^ Cranial window implantation in mice was done in a modification of what we had previously described, including a custom-made teflon ring for painless head fixation during imaging. 1 to 3 weeks after cranial window implantation, 50.000-100.000 glioblastoma cells were stereotactically injected into the mouse cortex at an approximate depth of 500 μm. Alternatively, the stereotactic tumor injection was performed without prior cranial window implantation into the mouse cortex as described above or into the striatum (1 mm anterior to bregma and 2 mm lateral to midline, 2 mm deep to cranial surface).

For *in vivo* retrograde tracing of the neuronal connectome, tumor injections were done following the direct labeling protocol as described above. Tumor cells were injected either into the cortex or the striatum. For *ex vivo* analyses of tissue, mice were sacrificed via perfusion between 14-30 days following tumor implantation.

#### Intravital microscopy

For *in vivo* two-photon imaging, male NMRI nude mice were implanted with cranial window and injected with tumor cells as described previously.^9,16^ The tumors were observed from 1 week after tumor implantation with a Zeiss 7MP setup (Zeiss) equipped with bandpass filter sets of 500 - 550 nm and 575 - 610 nm, using a 20x (1.0 NA) apochromatic, 1.7 mm working distance, water immersion objective (Zeiss). A pulsed Ti:Sapphire laser (Chameleon II ultra; Coherent) was used at 960 nm wavelength.

Isoflurane gas was diluted in 100% O2 to a concentration between 0.5 - 2.0% for *in vivo* imaging. For the induction of anesthesia, the mice were exposed to 4% isoflurane, which was lowered to 0.5-2% for the rest of the experiment and was monitored throughout the experiment. Eye cream was applied after anesthesia induction. During imaging, the body temperature was monitored and kept at 37°C using a temperature sensor and a heating plate. Anesthesia was regularly evaluated during image acquisition by checking the breathing rate.

Stacks from each time point of dendritic plasticity time-lapse imaging were hyperstacked and registered in Fiji by using a custom script.^108^

#### Intravital microscopy analysis

Analysis of time-lapse imaging of dendritic plasticity in connected^TUM^ neurons was performed manually. After registration of each stack to minimize drift between acquisition time-points, regions of interest of dendritic stretches were cropped for further analysis. For each time-point, the number of dendritic spines from five total dendritic stretches was determined.

#### Sample preparation, immunohistochemistry, in situ, and confocal microscopy

For *ex vivo* analyses of PDX models, the mice were anesthetized with either ketamine/xylazine or pentobarbital i.p. First, mice were perfused transcardially with PBS followed by 4% PFA (w/v) in 1x PBS. After removal of the brain, it was post-fixed in 4% PFA overnight and kept in PBS at 4°C. Serial sections of 80-100 μm were cut with a semiautomatic vibratome (Leica VT1000s). For *in vitro* analyses, coverslips were washed once with 1x PBS and subsequently fixed with 4% PFA (w/v) in 1x PBS for 5-10 minutes. Afterwards, they were washed once with 1x PBS and stored in PBS at 4°C.

*Ex vivo* mouse brain slices and organotypic slices were first permeabilized with 5% (v/v) FBS and 1% (v/v) Triton X-100 in 1x PBS for 2 hours. In the following, the primary antibodies were solved in in 1% (v/v) FBS and 0.2% (v/v) Triton X-100 in 1x PBS with a general dilution of 1:100, with the exception of anti-Nestin mouse with a dilution of 1:300 and anti-GFP chicken with a dilution of 1:300. Afterwards, the slices were washed 3x with 2% (v/v) FBS in 1x PBS for 15 minutes each. The secondary antibodies were solved in the same buffer as the primary antibodies with a general dilution of 1:500. The primary and the secondary antibodies were both incubated for 20-24 hours each. After the incubation time of the secondary antibody, the slices were washed 3x with 1% (v/v) FBS in PBS for 10 minutes each, followed by 3x washing steps with 1x PBS for 10 minutes each. All incubation steps were performed at room temperature on a shaker. Sample mounting was performed with “SlowFade Gold” solution.

For *in vitro* stainings, the coverslips were permeabilized for 10 minutes with 0.2% (v/v) Triton X-100 in 1x PBS. Afterwards, blocking was performed by incubating the samples in 10% FBS (v/v) in 1x PBS for 10 minutes. In general, the primary antibodies were solved in blocking buffer with a dilution of 1:100, with the exceptions of anti-Nestin mouse with a dilution of 1:300 and anti-GFP chicken with a dilution of 1:200-300. Subsequently, after 1h of incubation, the coverslips were washed 2x with 1x PBS for 5 minutes each before the respective secondary antibody was applied with a general dilution of 1:500 in the blocking buffer. After another hour of incubation, the coverslips were washed again 2x with 1x PBS for 5 minutes each. All incubation steps were performed at room temperature, shaking. Finally, the coverslips were mounted with “SlowFade Gold” solution and DAPI diluted 1:10000 (v/v) in 1x PBS.

Images were acquired using either a 20x air (NA 0.8) or 63x oil immersion objective (NA 1.4) at a confocal laser-scanning microscope (LSM710 ConfoCor3 or LSM980 Airyscan NIR, Zeiss).

#### Airyscan microscopy of *ex vivo* brain slices

Airyscan microscopy of dendritic stretches of connected^TUM^ neurons was performed using LSM980 Airyscan NIR (Zeiss) with a 63x oil immersion objective (NA 1.4). Images were acquired using calibrated Airyscan detectors with a lateral resolution of 0.043 μm/pixel and an axial resolution of 0.15 μm/pixel. Airyscan processing was performed in the Zen Blue software.

#### Mouse and rat cortical, hippocampal and basal forebrain cultures

Preparation of rat cortical cultures was done as described previously.^16^ Briefly, cells from E19 embryos were seeded on 12 mm coverslips in 24-well plates coated with poly-L-lysine at a density of 90,000 cells/cm^2^. They were cultured in a medium of Neurobasal (Invitrogen), supplemented with B27 (50x, 2%v/v) and L-glutamine (0.5 mM). The same protocol was used for rat hippocampal cultures, with the exception of 2.5% Trypsin (10x) instead of 0.05% Trypsin-EDTA (1x), as used for cortical cultures.

Mouse cortical cultures were prepared similarly to rat cortical cultures using cells from P1 and P2 mouse pups.

Primary basal forebrain cultures were prepared as previously described from the dissected septum of E19 rat embryos and plated at a density of 100,000 - 200,000 cells per well on 12 mm coverslips in 24-well plates coated with poly-L-lysine.^109^ They were cultured in neurobasal medium supplemented with B27 supplement (50x, 2% v/v), L-glutamine (0.5 mM) and neuronal growth factor (50 ng/ml). Culture medium was changed twice a week.

#### Human iPSC- and ESC-derived neurons

Human embryonic stem cells (hESC) of line WA01/H1 were obtained from WiCell whereas iPSCs were locally derived from a healthy donor (HD6, Heidelberg University, Heidelberg, Germany). Pluripotent cells were feeder-free cultured on Matrigel-coated (Corning #15505739) dishes, using mTeSR Plus medium (StemCell Technologies #100-0276). mTeSR was changed every other day and cells were passaged every 3–5 days using ReLeaSR (StemCell Technologies #05872). All cell cultures were maintained in a humidified incubator with 5% CO2 at 37°C. All procedures were approved by the Robert Koch Institute.

Induced glutamatergic neurons were differentiated from iPSCs or hESC according to previously described methods.^54^ Briefly, for each differentiation 250,000 hESCs were detached with Accutase (Gibco), plated on matrigel-coated wells in mTeSR Plus containing Rho kinase inhibitor (Y27632, Axon Medchem #1683, or Thiazovivin) and simultaneously transduced with lentiviruses FU-M2rtTA and Tet-O-Ngn2-puromycin. One day later (defined as DIV0), the media was replaced with N2 media [DMEM/F12 (Gibco #11330032), 1% N2 supplement (Gibco 17502048) 1% non-essential amino acids (Gibco #11140050), laminin (200 ng/ml, Thermo Fisher #23017015), BDNF (10 ng/ml, Peprotech #450-02) and NT-3 (10 ng/ml, Peprotech #450-03) supplemented with Doxycycline (2 μg/ml, Alfa Aesar) to induce expression of Ngn2 and the puromycin resistance cassette. On DIV1, puromycin (1 mg/ml) was added to the medium and after 48h of selection, cells were detached with Accutase (Gibco #A1110501) and re-plated on Matrigel-coated coverslips along with mouse glia (see paragraph below, typically at a density of 150,000 iGluts/24-well) in B27 media [Neurobasal-A (Gibco #12349015 supplemented with B27 (Gibco #17504044), GlutaMAX (Gibco #35050061) laminin, BDNF and NT-3]. Near 50% of the medium was replaced every second day for eight days, with cytosine arabinoside (ara-C; Sigma #C6645) added to a working concentration of 2 μM to prevent glia overgrowth. From DIV10 onward, neuronal growth media [Neurobasal-A supplemented with B27, GlutaMAX and 5% fetal bovine serum (FBS) (Hyclone #SH30071.03HI)] was washed in and used for partial media replacements every 3-4 days until analysis, typically after 4-6 weeks in culture.

Mouse glia cells used for co-cultures with induced glutamatergic neurons, were isolated as described before.^110^ Briefly, P3 mouse cortices from wildtype C57BL6 mice were dissected and triturated with fire polished Pasteur pipettes, and passed through a cell strainer. Typically, lysates from two cortices were plated onto a T75 flask pre-coated with poly-L-lysine (5 mg/ml, Sigma #P1274) in DMEM supplemented with 10% FBS (Sigma). Once primary mouse glial cells reached confluence, they were dissociated by trypsinization and re-seeded twice and then used for co-culture with induced glutamatergic neurons.

#### Cell viability assays

To assess toxicity of rabies virus to patient-derived glioblastoma spheroids, cells were seeded on to an opaque 96 well plates in neurobasal medium supplemented with B27 (50x, 2% v/v) and L-glutamine (0.5 mM), at a density of 5000 cells/well. Per patient-derived glioblastoma model, we seeded wells with glioblastoma cells transduced only with the TVA-oG-mCherry construct, directly labeled TVA-oG-mCherry and CVS-N2c^ΔG^-eGFP(EnVA) expressing glioblastoma cells, and directly labeled TVA-oG-mCherry and SAD-B19^ΔG^-eGFP(EnVA) expressing glioblastoma cells, including a control with only medium to measure background signal. The assay was performed according to the manufacturers protocol (Promega, Madison, WI) after 24, 48 and 72 hours. Luminescence was measured 10 minutes after incubation at room temperature for signal stabilization.

#### Direct and sequential labeling of glioblastoma cells for retrograde tracing

Experiments were performed following either the direct or the sequential labeling approach. For the direct approach, patient-derived glioblastoma spheroids were transduced with both the TVA-oG-mCherry construct and either SAD-B19^ΔG^-eGFP(EnVA) (5x10^4^ vg/ml) or CVS-N2C^ΔG^-eGFP(EnVA) (10^6^ vg/ml) used in this study before conducting further experiments. They were cultured as described above under spheroid primary culture conditions. For the sequential approach, TVA-oG-mCherry expressing glioblastoma cells were seeded, followed by a sequential rabies infection on the co-cultures at a titer depending on paradigm.

#### Sparse and dense sequential retrograde labeling

For tracing of the neuronal connectome of singular tumor cells, SAD-B19^ΔG^-eGFP(EnVA) or CVS-N2C^ΔG^-eGFP(EnVA) were added to TVA-oG-mCherry seeded co-cultures 2 hours after seeding on DIV07 rat cortical neurons at a titer of 10 vg/ml. For dense labeling of glioblastoma cells, SAD-B19^ΔG^-eGFP(EnVA) or CVS-N2C^ΔG^-eGFP(EnVA) were applied at a titer of 10^5^ vg/ml.

#### *In vitro* live cell time-lapse imaging of retrograde labeling

For rabies virus based retrograde live cell imaging, TVA-oG-mCherry expressing patient-derived glioblastoma spheroids were seeded onto DIV07 rat cortical cultures at a density of 1000 cells per well in 24 well plates. SAD-B19^ΔG^-eGFP(EnVA) or CVS-N2C^ΔG^-eGFP(EnVA) (both 10^3^ vg/ml) virus was added 1 hour after seeding. For experiments at later infection time points, rabies viruses were added 5 or 11 days after seeding glioblastoma cells.

Imaging was performed 2 hours after seeding of glioblastoma cells for a time period of 3-5 days at 37 degrees Celsius with 5% CO2. Images were acquired using a Zeiss LSM780/710 Zeiss Celldiscoverer7 confocal or a Nikon Ti-HCS widefield microscope with a 10x (NA 0.3)/20x (NA 0.95) objective and a pixel size of 770nm – 1.38µm. Coverslips were scanned every 20-45 minutes.

#### *In vitro* live cell time-lapse imaging of neuron-tumor co-cultures

For live cell experiments, tdTomato or GFP transduced patient-derived glioblastoma cells were seeded onto DIV7 rat cortical cultures at 1000 cells per well. For glioblastoma monocultures, 1000 cells per well were seeded in 24 well plates containing the same medium as co-cultures, namely Neurobasal (Invitrogen) supplemented with B27 (50x, 2% v/v) and L-glutamine (0.5 mM). Co- and monocultures were imaged at the same DIV, 4-13 days after seeding. Patient-derived glioblastoma cells were imaged for a period of 12-18 hours at 37 degrees Celsius with 5% CO2. Images were acquired using a Zeiss LSM 780 confocal microscope with a fully open pinhole every 10 minutes, with a 10x (NA 0.3) air objective and a pixel size of 346nm.

#### Invasion Speed Analysis with Trackmate

Field of views of *in vitro* live cell imaging data were analyzed in Trackmate (version 7.11.1).^111^ The Kalman tracker was used (parameters: Initial search radius = 50, Search radius =30, Max frame gap = 10). For quality control, only tracks with track durations over 10000s were kept to account for false tracks made by the tracking algorithm.

#### Infection lag time analysis

For determination of approximate infection lag time, images from live cell time-lapse imaging of tumor cells infected at DIV00 were used. The time point of GB^Starter^ cell infection was manually determined when a TVA-oG-mCherry expressing tumor cell became visually eGFP-positive after rabies virus infection with SAD-B19^ΔG^-eGFP(EnvA) or CVS-N2c^ΔG^-eGFP(EnvA). Earliest connected^TUM^ neuron infection was calculated by subtracting time point of visible infection of first connected^TUM^ neuron in vicinity of infected GB^Starter^ from the time point of GB^Starter^ infection.

#### Drug treatment and radiotherapy in co-cultures

For drug treatment experiments, coverslips were treated with an end concentration of 40µM perampanel 2 hours post glioblastoma cell seeding. Controls were treated with respective amount of DMSO. Coverslips were imaged on the same day of seeding and then 3, 5 and 7 days after seeding using a Zeiss LSM 780 microscope with a 10x air (NA 0.3) objective at 37 degrees Celsius with 5% CO2.

For irradiation experiments, glioblastoma cells were seeded on to DIV7 rat cortical neurons in 24-well plates (1000 cells/well). For combined perampanel treatment and radiotherapy, coverslips were treated with 40 µM perampanel 2 hours after seeding. 5 days after seeding tumor cells, coverslips were irradiated at 4 Gray. For radiotherapy in combination with retrograde labeling of patient-derived glioblastoma cells, irradiated and control coverslips were infected with CVS-N2c^ΔG^-eGFP(EnVA) virus (10^3^ vg/ml) 6 hours after irradiation. Coverslips were fixed 3 days later and analyzed for input-to-starter ratios (see method section “determination of input-to-starter ratios”).

#### Rabies virus-based genetic ablation of connected^TUM^ neurons

DIV06 rat cortical neurons were infected with AAV5 virus based on the AAV-Flex-TACasp3-TEVP plasmid (Addgene #45580)^83^ at a titer of >7x10^8^ vg/ml. AAV-flex-taCasp3-TEVp was a gift from Nirao Shah & Jim Wells (Addgene plasmid #45580; http://n2t.net/addgene:45580; RRID: Addgene_45580). The following day, all wells were washed 3x with pre-warmed culture medium (Neurobasal with B27 (50x, 2% v/v) and L-glutamine (0.5mM) before seeding 1000 TVA-oG-mCherry expressing glioblastoma cells per well. 2 hours after seeding, SAD-B19^ΔG^-Cre-GFP(EnVA) (based on Addgene plasmid #32634) was added at a titer of 10^4^ vg/ml.^112^ Control wells were treated with the same concentration of SAD-B19^ΔG^-Cre-GFP(EnVA) but without prior infection of neuronal cultures with AAV-Flex-TACasp3-TEVP. 10 days after seeding of tumor cells, coverslips were fixed and stained with human-specific anti-Nestin (Abcam, 22035) to label glioblastoma cells as previously described.^8,16^ Quantification was done as described in methods section “determination of input-to-starter ratios”.

#### Single-cell RNA sequencing

For single-cell RNA sequencing of rabies transduced cultures and their controls, co-cultures of rat cortical cultures and human glioblastoma cells were processed on DIV06. First, the cells were dissociated from coverslips by incubating with Trypsin for 5 minutes. Then, the cells were collected in falcon tubes and centrifuged before resuspending in FACS buffer (10% FBS in PBS). DAPI was used at a final concentration of 1 µg/ml as a cell viability marker. Sorting was performed with FACSymphony S6 (BD Biosciences). GB^Starter^ were identified by simultaneous GFP and mCherry fluorescence. Connected^TUM^ neurons were identified by the GFP signal and cells without fluorescence signal were categorized as unconnected^TUM^ microenvironmental cells. The following filters were used: 450/20 for DAPI, 530/30 for GFP and 610/20 for mCherry. Lasers with wavelengths of 405 nm, 488 nm and 561 nm were used for this purpose.

#### Sequencing pre-processing and analysis

The analysis of the single-cell RNA sequencing data was performed using the R package Seurat (version 5.0.1)^113^ unless indicated otherwise. The sequencing data was preprocessed and high-quality rat cells matching the following criteria were analyzed: unique number of transcripts (5,000-11,250), number of reads (100,000-2,000,000), fraction of mitochondrial reads less than 4%. The number of highly variable features was set to 4,000 and data integration was performed using the Seurat method “CCAIntegration”. The connected^TUM^ neurons were identified based on the eGFP expression level as measured by FACS.

##### Identification of cell types

Previously published gene sets were used to identify different cell types and states (annotation level 3).^114^ To this end, the expression of a gene set across the clusters was assessed using the Seurat module score function. Astrocytes and oligodendrocytes were identified by a mean module score > 0.1 in the respective gene set, which was in line with the expression of known marker genes. The subanalysis of neurons was performed after excluding astrocytes and oligodendrocytes from the dataset.

#### Invasivity module score for single-cell RNA sequencing analyses

Invasivity scores for the different patient-derived glioblastoma models were calculated as described before.^8^ Briefly, pseudotime was estimated, with initial cells being designated as SR101-negative invasive cells. Genes exhibiting either positive or negative correlation to pseudotime across cell lines were identified. Following this, the invasivity score was determined by subtracting the Module score of genes negatively correlated from that of positively correlated genes.

#### Synaptogenic module score for single-cell RNA sequencing analyses

The “Synapse assembly” GO term was downloaded from https://amigo.geneontology.org/amigo/.^61,62^ Synaptogenic score was calculated from the list of 117 genes using the AddModuleScore function from Seurat. Score correlations were calculated in r.

#### Single-cell neurotransmitter genes expression analysis

Using the AddModuleScore function from Seurat, a score was calculated for each neurotransmitter group of interest. Genes included for each group can be found in Table S2.

#### Analysis of publicly available singe-cell RNA sequencing data

Publicly available single-cell RNA sequencing data from various publications were used for analysis.^5,63,114^

#### Correlative *in vivo* two-photon and *ex vivo* volume electron microscopy

For conducting *in vivo* correlative light and electron microscopy (CLEM) with infrared branding, we detected the glioblastoma cells based on their mGFP expression and classified them based on their uptake of Sulforhodamine 101 (SR101), which was monitored with two-photon microscopy in living mice over time. After identification of the glioblastoma cells, the mice underwent transcardial perfusion following a previously established protocol.^8^

To facilitate the correlation process on a macroscopic level as well as on *ex vivo* and electron microscopy imaging level, approximately 7 ml of 2% Evans Blue were added into the last 20 ml of 4% PFA at the end of the perfusion. This addition was designed to enhance blood vessel signal *ex vivo*. To preserve the *in vivo* orientation for imaging, the mice were decapitated, leaving the window and the titanium ring on the head, allowing immediate subsequent two-photon microscopy with consistent positioning. For the infrared branding, cells of interest that were chosen before were centered in a field of view of 694 x 694 µm using a 16x objective. A high-resolution z-stack (pixel-size: 0.67 µm) of the area prior to branding was obtained.

To precisely localize the region of interest within the brain *ex vivo*, an infrared branding was performed.^115^

This macroscopically visible region, providing clear demarcation for the region of interest, was cut out of the brain in the form of a cube using a surgery knife. Subsequently, the cube was embedded in agarose in a manner that its surface was parallel to the sectioning blade of the vibratome. Slices of 300 μm thickness were obtained using a Leica Vibratome.

The sample was then stained with DAPI to provide further landmarks for correlative imaging. Preparation of the sample for electron microscopy was performed as previously described.^8,16,116^ Then, we captured low-resolution overviews with serial-section scanning electron microscopy. The cells of interest were identified by their cell morphology, by their spatial arrangement of the DAPI-stained nuclei and the Evans blue signal as a marker for blood vessels. Lastly, we acquired the cells of interest throughout large z-volumes and reconstructed them.

Stable glioblastoma cells were discerned by longitudinal, intravital imaging and through the colocalization of SR101 with the intrinsic mGFP signal exhibited by the tumor cells.

#### Correlative *ex vivo* confocal microscopy and *ex vivo* volume electron microscopy

We followed the sample preparation, microscopy and analysis protocols for *ex vivo* correlative light and scanning electron as described before.^8^ The mice were administered anesthesia using pentorbital i.p. We initiated the perfusion process with transcardial infusion of PBS, succeeded by a 4% solution of PFA in PBS. After extracting the mouse brain, it was subjected to an overnight post-fixation in 4% PFA and subsequently preserved in PBS at 4°C for storage. We prepared serial brain sections of 80 µm thickness using a semi-automatic vibratome (Leica VT1000s) which were then screened under a widefield fluorescence microscope (Leica DM6000) to check for intrinsic tumor cell fluorescence. Subsequently, we carefully cut tissue blocks with an approximate area of 400 μm^2^ for in-depth analysis and volume electron microscopy.

Our approach relied on CLEM for the identification of cells of interest. Therefore, the sections were stained with DAPI (1:10000) and subsequently imaged under a Leica TCS SP8 confocal microscope (63x objective [NA 1.4] or 20x objective [NA 0.75]; pixel size 200 nm; z-stack with 520 nm-steps; scanning speed of 400-600 Hz). The rinsing-, post-fixation-, and contrasting-steps for the electron microscopy processing were conducted as described before.^8,116,117^

#### Electron microscopy and image analysis

Electron microscopic imaging was performed using a LEO Gemini 1530 scanning electron microscope (Zeiss) and an Auriga scanning electron microscope (Zeiss) in combination with an ATLAS scan generator (Zeiss). Imaging and synapse identification was performed as described previously.^8^

To visualize the distribution of the synaptic vesicles and their distance to the synaptic cleft, the “scatterplot3d” function in R^118^ was used. The vesicles were plotted in three dimensions with color indicating the distance based on calculations on the 2D-sections.

The cell and synaptic bouton boundaries were manually segmented and imported as area-lists on consecutive EM sections.^119^ 3D representations and renderings of the synaptic inputs on stable and invasive glioblastoma cells were performed in Arivis Vision4D x64 and 3dmod. During this visualization, mesh surfaces were refined. Subsequently, the exported video renderings were montaged in DaVinci Resolve.

#### Determination of input-to-starter ratios

Patient-derived glioblastoma spheroids were seeded onto DIV7 rat cortical neurons following either the direct or the sequential labeling protocol as described above. 8 days later, coverslips were fixed and stained for Nestin and GFP as described above.

For input-to-starter ratio analysis of highly invasive versus non-invasive regions, coverslips were infected with CVS-N2c^ΔG^-eGFP(EnvA) (10^3^ vg/ml) at DIV05 (highly invasive) or DIV11 (non-invasive) and fixed 3 days after virus application. Coverslips were imaged at a Zeiss AxioScanZ1 microscope with a 20x (NA 0.8) objective and a pixel size of 325 nm.

Cell somata were trained using the ilastik^120^ pixel classification pipeline. Probability maps were exported. All further processing steps were performed in Fiji.^108^ For quantification of the number of tumor cells per coverslip, probability maps of Nestin and DAPI signals were multiplied. Afterwards, the resulting image was auto-thresholded using the “Threshold” function and the number of cells were then determined using the “Analyze Particles” function with a cut-off by a minimum of 20 µm diameter. Auto-thresholding and particle analysis was also performed for the probability map of the eGFP channel. To extract the number of GB^Starter^ cells, thresholded images were multiplied and resulting particles were counted. The number of input cells (connected^TUM^ neurons) were calculated by subtracting the number of GB^Starter^ cells from the number of all eGFP-positive cells.

#### Cell type analysis of connected^TUM^ neurons

For *ex vivo* analysis, brain slices obtained from mice sacrificed 14-30 days after tumor injection with TVA-oG-mCherry expressing, SAD-B19^ΔG^-eGFP(EnvA) or CVS-N2c^ΔG^-eGFP(EnvA)- infected cells from patient-derived glioblastoma spheroids were stained for Nestin or mCherry, GFP and a marker of interest from the above listed. Slices were imaged at a Leica DM6000 microscope with a 10x (NA 0.4) objective.

Crops were manually analyzed for number of eGFP-positive cells, number of marker of interest positive cells and double positive cells.

For the *in vitro* quantification, TVA-oG-mCherry expressing, SAD-B19^ΔG^-eGFP(EnvA) or CVS-N2c^ΔG^-eGFP(EnvA)-infected cells from patient-derived glioblastoma spheroids were seeded onto DIV7 rat cortical neurons. 8 days later, coverslips were fixed and stained for Nestin, GFP and a marker of interest (used in our study were: NeuN, S100B, MBP, Iba1, CAMK2, Parvalbumin, Chat). Coverslips were imaged at a Zeiss AxioScanZ1 microscope with a 20x (NA 0.8) air objective and a pixel size of 325nm.

For quantification of DLX-infected connected^TUM^ neurons, DIV6 rat cortical neurons were treated with AAV-mDlx-NLS-mRuby (Addgene #99130) at a titer of >1x10^9^ vg/ml.^121^ AAV-mDlx-NLS-mRuby2 was a gift from Viviana Gradinaru (Addgene plasmid #99130; http://n2t.net/addgene:99130; RRID: Addgene_99130). One day later, glioblastoma cells infected with the direct labeling approach were seeded at 1000 cells/well. Coverslips were fixed at tumor cell div08 and quantified as described above.

To rule out unspecific leakage or labeling of rabies virus we have done different control experiments. First, we have exchanged medium of wells with TVA-oG-mCherry expressing and SAD-B19^ΔG^-eGFP(EnvA) or CVS-N2c^ΔG^-eGFP(EnvA)-infected tumor cell co-cultures of both strains onto wells of only rat cortical neurons and fixed these 8 days after medium exchange. Furthermore, we have seeded lysed TVA-expressing, rabies-infected glioblastoma cells onto DIV7 rat cortical neurons. Cells were lysed by first exposing them to sterile water and subsequently mechanically lysing them by pipetting them through a 25-gauge needle for 60 seconds. Cell lysis was confirmed by Trypan-blue staining when counting cells.

#### 4ExM Expansion microscopy

4x expansion microscopy was performed as described before.^122^ Briefly, mouse cortical and iPSC co-cultures with human glioblastoma cells were cultured as described above. Immunohistochemistry was performed as described to stain against GFP and Nestin, since endogenous fluorescence is expected to be quenched after expansion. Following immunohistochemistry, coverslips were anchored in 0.1 mg/ml Acryloyl-X SE (AcX) solution in 1x PBS overnight at room temperature. AcX stock solution was 10 mg/ml AcX solved in DMSO. Coverslips were then incubated for 2 hours at 37° C in custom chambers in 100 µl of the gelation solution consisting of 470 µl monomer stock solution for 4xM (0.08 % (v/v) sodium-acrylate (33% wt stock), 2.5% (v/v) acrylamide (50% wt stock), 0.02% (v/v) cross-linker (1% wt stock), 1.9M NaCl (5M stock), 1 ml of 10x PBS, 18.8% (v/v) water) mixed with 10 µl each of 0.5 wt% 4-HT, 10 wt% TEMED and 10 wt% APS. All stock solutions were prepared with water. Incubation chambers were prepared using microscope slides and spacers made from No. 0 coverslips. After incubation, gels were recovered and digested in 8 U/ml Protein-K buffer overnight at room temperature. The gels were then stained with DAPI (1 µg/ml in 1x PBS) for 30 min and washed afterwards for 30 min with 1x PBS. The gels were expanded by washing with MilliQ water 3x 10 min followed by 1x 30 min at room temperature.

Expanded gels were mounted on and imaged in poly-L-lysin coated glass bottom dishes.

The scale bars in expansion microscopy images shown in figures were placed after accounting for the expansion factor of 4.

#### Tissue clearing

Whole brain immunolabeling was performed according to the iDISCO+ protocol.^123^ Briefly, samples were dehydrated with a methanol/PBS series (catalog # 8388.2; Carl Roth, Karlsruhe, Germany): 20 vol%, 40 vol%, 60 vol%, 80 vol%, 100 vol% (twice) for one hour each, followed by overnight incubation in 66 vol% Dichloromethane (DCM) (KK47.1, Carl Roth) and 33 vol% methanol. Samples were then washed twice in 100 vol% methanol followed by a bleaching step with 5% H_2_O_2_ (catalog # LC-4458, Labochem, Sant’Agata li Battiati, Italy) overnight at 4 °C. Rehydration was performed with a methanol/PBS series containing 80 vol%, 60 vol%, 40 vol%, 20 vol%, PBS for 1h each. Lastly samples were washed in 0,2 vol% TritonX-100 (x100, Sigma) in PBS (PTx.2) twice for 1h.

Immunolabeling was performed by incubating pretreated samples in permeabilization solution (400 ml PTx.2, 11.5 g Glycine (catalog # G7126, Sigma), 100 ml DMSO (catalog # A994, Carl Roth) for 2 days at 37° C. Brains were then blocked in blocking solution (42 ml PTx.2, 3 ml Donkey serum, 5 ml DMSO) for 2 days at 37° C. For immunolabeling primary antibodies for against GFP (catalog # GFP-1020, Aves Labs) and against RFP (catalog # 600-401-379, Rockland) were applied in PBS, 0,2% Tween-20 (P2287, Sigma) (PTw), 5% DMSO, 3% goat serum for 7 days at 37° C on a rocking platform. Then samples were washed in PTw for 5 times until the next day and incubated with secondary antibodies (goat anti-chicken 488, catalog # A32931, Thermo Fischer and goat anti-rabbit 568, catalog # A11011, Thermo Fischer) in PTw and 3% goat serum for 7 days at 37° C. Samples were wrapped in aluminum foil to prevent photobleaching. Samples were washed in PTw for 5 times until the next day.

Clearing was performed by dehydrating the samples in a methanol/PBS series: 20 vol%, 40 vol%, 60 vol%, 80 vol%, 100 vol% (twice) for one hour each. Followed by 3h incubation in 66 vol% DCM and 33 vol% Methanol, samples were incubated twice in 100 vol% DCM for 15 minutes. Lastly samples were incubated in 33 vol% benzyl alcohol (catalog # 24122, Sigma) and 67 vol% benzyl benzoate (vol/vol; catalog # W213802, Sigma) without shaking.

Unless otherwise stated all steps were performed at room temperature, while shaking. Clearing agents were freshly prepared for each step of the protocol.

#### Light-sheet microscopy

Cleared samples were imaged with a light-sheet microscope (Ultramicroscope II, Miltenyi Biotec, Heidelberg, Germany) using a 4x objective (MI Plan objective lens 4x, NA 0.35) and combined lasers (excitation wavelength at 470 nm and 560 nm). The in-plane resolution was 1.63 x 1.63 µm with a step size of 5 µm. Images were stitched with a custom-made macro in Fiji/ImageJ.^124^

#### Calcium imaging of connected^TUM^ and unconnected^TUM^ neurons

6 days after seeding of rat cortical neurons, cultures were infected with AAV.Syn.NES-jRGECO1a.WPRE.SV40 (Addgene #100854).^125^ AAV.Syn.NES-jRGECO1a.WPRE.SV40 was a gift from Douglas Kim & GENIE Project (Addgene plasmid #100854; http://n2t.net/addgene:100854; RRID: Addgene_100854). The following day, TVA-oG-mCherry expressing, CVS-N2c^ΔG^-eGFP(EnvA)-infected glioblastoma cells were seeded at a density of 1000 cells/well. 12 days after seeding, cultures were imaged on a Zeiss LSM 980 confocal microscope with a 20x objective (NA 0.8) with a pixel size of 345.26 nm and a frame interval of 0.52 sec.

#### Calcium analysis of connected^TUM^ and unconnected^TUM^ neurons

For the analysis of calcium transients, somata of connected^TUM^ and unconnected^TUM^ neurons were marked with circular regions of interest. Mean gray value and center of mass were multi-measured in Fiji^108^ for all imaging time-points. The exported measurements were further quantified using a custom-written MATLAB script.^8^

#### Functional neurotransmitter receptor screening

Calcium imaging experiments were performed with a triggered neurotransmitter puffing onto the glioblastoma cells. They puffing pipettes were placed approximately 30μm above the targeted region of interest (ROI). For these recordings, Patchmaster software (HEKA) was used, with a puff applied every 45 s. Puffing stimulations were generated at 10-15 PSI using a Picospritzer. Each recording lasted 225 seconds. Images were acquired with pixel sizes of 1,3 μm, 0.2 μm and 0.7 μm at a Leica TCS SP5 microscope using a 20x (NA 0.5) water objective, respectively. The recoding frequency was 1.56 Hz in a bidirectional acquisition mode.

Pipettes for puffing were fabricated from borosilicate capillaries (World Precision Instruments) and had resistances of 2-7 MΩ. The pipettes were filled with 200 μl of the neurotransmitter stock and 0.4 μl of Alexa 594 coloring agent from Invitrogen.

Functional neurotransmitter receptor screening occurred by sequentially puffing 8 different neurotransmitters onto a region of interest to determine which trigger a response in glioblastoma cells. A baseline recording with aCSF puffing was used first to exclude regions with a non-neurotransmitter specific response. Next, glutamate puffing was performed for 225 seconds and 5 puff stimulations. Further, acetylcholine (1 mM), GABA (100 mM), ATP (1 mM), serotonin (5HT) (1.5 mM), adrenaline (1 mM), glycine (2 mM) and dopamine (10 mM) puffing followed under the same conditions. All neurotransmitter stocks were prepared with calcium-free aCSF and for puffing, 200 μl of aCSF with 0.4 μl Alexa 594 were used to visualize the neurotransmitter puff as control for successful neurotransmitter application during calcium imaging.

For pharmacological experiments, two baseline recordings with neurotransmitter- and control puffing were performed as described. Next, atropine (50nM) was washed in for 450 seconds to ensure that the coverslip was fully submerged, and a pharmacological effect could be observed. After the wash in, a third recording took place with Acetylcholine puffing under altered conditions. Atropine was then washed out with regular aCSF for 450 seconds. Finally, one last calcium imaging time-lapse recording was performed to assess whether the initial response of the cell could be recovered after washing out atropine.

#### Calcium imaging analysis

AQuA was used to quantify the event frequency, area, duration, ΔF/F and total calcium entering the cell (Area Under Curve) for each calcium event.^126^ The raw calcium imaging recordings contain two channels, one with the puff recording, and the other with the calcium signal. For the semi-automatic AQuA data analysis, the channels were split using a Fiji macro and the recording with the calcium signal was analyzed further. Single cells from each ROI were defined in a user interface and all cells were batch processed using the same detection settings for all files.

#### Whole-cell patch-clamp electrophysiology

Whole-cell patch clamp recordings were made from coverslips secured under a platinum ring in the recording chamber (OAC-1; Science Products) and submerged in continuously flowing (3 mL/min) artificial cerebrospinal fluid (aCSF, in mM: NaCl, 125; KCl, 3.5; CaCl_2_, 2.4; MgCl_2_, 1.3; NaH_2_PO_4_, 1.2; glucose, 25; NaHCO_3_, 26; gassed with 96% O_2_ and 4% CO_2_) maintained at 32–34 °C with an in-line perfusion heater (TC324B; Warner Instruments). Patch electrodes (3-5 MΩ) were pulled from 1.5 mm borosilicate glass. For paired recordings, action potential recordings and postsynaptic current recordings, the following internal solution was used (in mM): KMethylsulphate, 135; EGTA, 0.2; HEPES, 10; KCl, 12; NaCl, 8; Mg-ATP, 2; Na_3_-GTP, 0.3. Methylsulphate was used instead of gluconate as the principle intracellular anion to avoid a rundown of both sAHP amplitude and AP accommodation. Data were not corrected for the liquid junction potential of 10.1 mV calculated with JPCalc (RRID:SCR_025044). Recordings were made with a Multiclamp 700B amplifier, digitized through a Digidata 1550B A/D converter and acquired and analyzed using pClamp 11 software (Molecular Devices). Recordings commenced only after passive properties had stabilized and these values were used for analysis. Cells with an access resistance above 25 MΩ were excluded from analysis. Voltage clamp recordings were sampled at 20 kHz with a low pass filter of 2 kHz. Current clamp recordings were sampled at 250 kHz with a low pass filter of 10 kHz. Pipette, but not whole cell capacitance, was compensated in all recordings. For Biotin filling, 0.3% Neurobiotin Tracer from Vector Laboratories was used.

#### Patch-clamp analysis

Action potential and postsynaptic current analysis was performed in Easy Electrophysiology (RRID:SCR_021190): Rheobase current (the minimal required current injection step needed to evoke an AP), AP threshold, AP amplitude, half width, afterhyperpolarization (AHP) potential amplitude and AHP delay to peak were assessed from the first AP evoked by 1 s depolarizing current injection steps applied in 10 pA increments from a potential of -70 mV maintained by constant current injection. Spontaneous APs or any AP coinciding with current injection onset were excluded from analysis. AP threshold was defined as the point where the first derivative of the voltage trace reached 20 mV/ms during the rising AP phase. AP and AHP amplitudes kinetics were calculated relative to this threshold and rise and decay times represent 10 to 90% of the threshold to peak interval. Input/output functions represent the frequency of APs generated over a depolarizing current injection step of 1 s versus the current injection amplitude (in pA).

Miniature postsynaptic currents (mPSCs) were recorded at -70 mV in the presence of TTX (0.5 M). Due to the more positive chloride reversal potential (-49 mV), both GABA_A_ receptor-mediated inhibitory mPSCs (mIPSCs) and AMPA receptor-mediated excitatory mPSCs (mEPSCs) were recorded as inward currents distinguishable by their decay times: Events with a decay time (defined as time between the peak and the point at which the event decayed to 37% (1/e)) up to 10 ms and a rise time from 0.5 to 5 ms were defined as mEPSCs, while events with a decay time longer than 12 ms and a rise time from 0.5 to 15 ms were defined as mIPSCs. Thresholds for decay times were established from recordings in the presence of the GABA_A_ receptor antagonist gabazine (Biotrend, 5 µM), or the AMPA-receptor blocker 2,3-dihydroxy-6-nitro-7-sulfamoyl-benzo[f]quinoxaline (NBQX, Hello Bio, 5 µM) (n = 4 cells each). Events were detected via template matching after filtering with a 2000 Hz Bessel low-pass filter while a minimum amplitude threshold of 5 pA was used to exclude noise (RMS noise was < 5 pA). All events were fit with a biexponential function and visually verified. Decay kinetics were fit with a single exponential function with the formula:

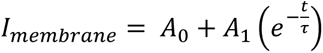

where I_membrane_, represents the membrane current, A_0_ and A_1_ represent the mean baseline current and slope parameter and τ the decay time constant.

Spontaneous network activity was evaluated from 3-5 min long current-clamp recordings without any holding current. EPSP bursts (>500 ms and >5 mV with multiple synaptic events) and APs were counted manually. Burst depolarization per second was calculated from the mathematical integral of the difference between the baseline membrane potential outside burst events (Savitzky-Golay smoothed) and the lower envelope of the EPSP burst after smoothing (500 point) to remove any APs. SIC and AP burst envelope decay kinetics were analyzed in cells with large and clean single peak responses and expressed as weighted tau values from biexponential fits using the following formula:

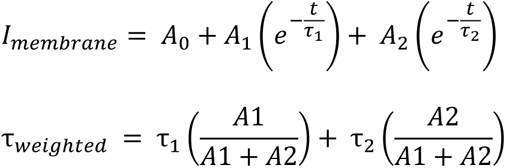

#### Electrophysiological characterization with high-density microelectrode arrays

Recordings were performed on multi-well high-density microelectrode arrays (HD-MEAs) available from MaxWell Biosystems (MaxTwo system, Zurich, Switzerland).^127^ Before the plating, HD-MEAs underwent sterilization using 70% ethanol for 30 minutes, followed by three successive rinses using distilled water. For enhanced tissue attachment, the arrays were treated with a coating of 0.05% poly(ethyleneimine) (Sigma-Aldrich), prepared in borate buffer at a pH of 8.5 (Thermo Fisher Scientific, Waltham, USA). This coating process was carried out for 30 minutes at room temperature. Subsequently, the arrays were rinsed again with distilled water and then allowed to air dry.

Embryonic day (E) 18 rat primary cortical neurons were prepared as described previously.^127^ Neurons were seeded at a density of 20-30’000 cells per chip in plating medium, which contained 450 mL Neurobasal (Invitrogen, Carlsbad, CA, United States), 50 mL horse serum (HyClone, Thermo Fisher Scientific), 1.25 mL Glutamax (Invitrogen), and 10 mL B-27 (Invitrogen). Primary cultures were housed in culture incubators at 37C/5% CO_2_. After two days, the plating medium was gradually changed to maintenance medium, which contained BrainPhys and SM1 (STEMCELL Technologies, Vancouver, #05792); ½ of the media was exchanged every 2–3 days. On day in vitro (DIV) 7, an activity scan was performed to screen for active electrodes on the HD-MEA and to select a suitable recording configuration for the tracking experiment. Up to 1024 read-out channels were selected based on the action potential amplitude values estimated during the activity scan. Next, tumor cells were dissociated and seeded onto the primary culture for co-culture. Starting from DIV7 onwards, co-cultures were recorded every 1–2 days until DIV12 with the same network recording configuration (recording duration: 30-60 mins). No media changes were performed during this period.

Results were obtained from a total of 4 controls and 8 neuron/tumor co-cultures, using multi-unit activity. The firing rate was estimated for all active channels (minimum firing rate: 0.05 Hz), and averaged over the full array. The bursts were detected on binned spike train activity (1 second bins), using an adaptive threshold based on the activity of each well (peaks above the mean + 1.5 standard deviation of the binned population activity).

#### Cluster analysis of connected^TUM^ neurons over time

In this methodology, we employed a systematic analysis of input and starter cells and their spatial relationships. First, GFP and Nestin signals were segmented using ilastik.^120^ Starter cells were calculated by overlapping the segmented GFP and Nestin signals. Input cells were identified by subtracting starter cells from the GFP signal. We extracted the coordinates of the input and starter cells. In further analysis, we utilized MATLAB to conduct clustering using the Density-Based Spatial Clustering of Applications with Noise (DBScan) algorithm. Parameters such as MinPts and epsilon were adjusted according to the characteristics of individual samples. The MATLAB script calculated distances between starter cells, input cells, and the resulting clusters. Moreover, it transformed the cluster boundaries into Regions of Interest (ROI) represented by Convex Hulls for enhanced delineation. Within each cluster, we evaluated the input-to-starter ratio to assess the distribution and composition of input cells relative to starter cells.

#### Whole brain atlas mapping of tumor cells and connected^TUM^ neurons

To register and analyze brain sections, we used the QUINT workflow consisting of three steps.^128^ First, the sections were registered to Allen Mouse Brain Common Coordinate Framework (CCF).^129^ Sections were then preprocessed and segmented for quantification.

##### Data Acquisition and Preparation

Brain sections from experimental mice were acquired using the Zeiss AxioScanZ1 microscope with a 20x (NA 0.8) objective. Sections were stained with DAPI prior to acquisition. The endogenous mCherry was used to identify glioblastoma cells and the connected^TUM^ neurons were identified by the endogenous GFP expression.

##### Image Registration and Processing

The aligned image series were registered to the atlas using QuickNII and VisuAlign tools^130^ to ensure accurate alignment across different brain sections. QuickMASK tool was utilized for generating masks corresponding to left-right hemisphere delineations.

##### Tumor and Connected^TUM^ Neuron Analysis

In order to define the distance from GFP-positive, connected^TUM^ neurons to the tumor mass, regions of interest (ROIs) were delineated where the mCherry signal was very dense. These ROIs were cleared from the GFP signal. DAPI and GFP signals were separately trained in ilastik^120^ to segment nuclei and connected^TUM^ neurons, respectively. To separate connected^TUM^ neurons, nuclei of connected^TUM^ neurons were calculated by overlapping the segmented GFP and DAPI channels. Additionally, the centroids of these nuclei were extracted.

##### Quantification and Visualization

The Nutil tool was utilized to quantify GFP-positive nuclei across different brain regions.^131^ Main tumor site and GFP-positive, connected^TUM^ neurons were visualized in 3D using MeshView^130^, providing insights into their spatial distribution and connectivity patterns.

##### Distance Determination and Plotting

Using the coordinates of each centroid of GFP-positive nuclei, the distances of connected^TUM^ neurons to the tumor mass were determined separating ipsilateral and contralateral hemispheres. Distances to tumor, differences across hemispheres, and cell distribution were quantified and visualized for each experimental group.

##### Magnet resonance imaging

MR scans were conducted using a 9.4 Tesla horizontal bore small animal MRI scanner (BioSpec 94/20 USR, Bruker BioSpin GmbH, Ettlingen, Germany) equipped with a gradient strength of 675 mT/m and a receive-only 4-channel surface array coil. T2 weighted images of *ex vivo* brain samples were acquired using a 3D TurboRARE sequence (TE: 78.9 ms, TR: 1800 ms, spatial resolution: 0.1 x 0.1 x 0.1 mm3, FOV: 15 x 20 x 10 mm3, matrix: 150 x 200 x 100, averages: 1, flip angle: 180°, RARE factor: 25, time of acquisition: 12min 0s).

##### General image processing and visualization

Image processing was primarily performed in ImageJ/Fiji (e.g. to reduce and remove unspecific background by subtraction of different channels, filtering with a median filter or the ‘Remove Outlier’).^108^

Arivis Vision 4D and ImageJ/Fiji were used for 3D and 4D image visualization. Probability maps were created for further analysis and visualization using ilastik.^120^ For all 3D renderings in Arivis Vision 4D, probability maps were used. Confocal Laser Scanning Microscopy (CLSM) images and *in vivo* imaging data were denoised using the denoise.ai pretrained model in the Nikon NIS-Elements AR software v5.30.01 (Nikon GmbH Germany/Laboratory Imaging). Videos were produced in DaVinci Resolve 17.

#### Quantification and statistical analysis

Quantification results were analyzed in GraphPad Prism (GraphPad Software) or R to test statistical significance with the respective tests. Data were first analyzed for normality using D’Agostino and Pearson normality. For normally distributed data, statistical significance was determined by using the two-sided Students’ t-test. In the case of non-normality, Mann-Whitney test was used. For > 2 groups, normally distributed data were analyzed using a one-way ANOVA test and non-normally distributed data were analyzed with a Kruskal-Wallis (unpaired) or Friedman (paired) test. If the p value was below 0.05, results were considered statistically significant. Manual quantifications were performed by two independent investigators. Animal group sizes were kept as low as possible. No statistical methods were used for predetermining sample size. Quantifications were depicted with mean and standard error of means.

## DATA AVAILABILITY

All sequencing data will be deposited to GEO prior to publication.

## CODE AVAILABILITY

Code used for analysis is available at https://github.com/venkataramani-lab/.

## ACKNOWLEDGMENTS

The work was supported by the Deutsche Forschungsgemeinschaft (DFG, German Research Foundation), SFB 1389, UNITE Glioblastoma, project ID 404521405 (addressed to V.V., M.B.), project number VE1373/2-1516 (addressed to V.V.), Heidelberg University and Research Seed Capital (RiSC) from the Ministry of Science, Research and the Arts Baden Württemberg (addressed to V.V.). S.K.T., E.R., M.C.S were supported by the Deutsche Krebshilfe/German Cancer Aid (Mildred-Scheel-Scholarship for MD students). We gratefully acknowledge the data storage service SDS@hd supported by the Ministry of Science, Research, and the Arts Baden-Württemberg (MWK). This publication was supported through state funds approved by the State Parliament of Baden-Württemberg for the Innovation Campus Health + Life Science Alliance Heidelberg/Mannheim. C.P.B. was supported by the German Research Foundation (DFG: BE7081/2-1). We are grateful to Hilmar Bading, Department of Neurobiology and Interdisciplinary Center for Neurosciences, Heidelberg University, for providing the opportunity to carry out electrophysiological characterization of neuron-tumor networks in his laboratory. Rabies viruses for initial experiments were a gift from Karl-Klaus Conzelmann. We thank the Viral Core Facility Charité for supplying viral constructs used in this study. We thank M. Kaiser, M. Schmitt, F. Gleiche, V. Buchert and K. Eghbalian for technical assistance. We thank Y. Dörflinger and S. Hoppe for technical assistance with electron microscopy. We thank K. Becker, K. Dell, A. Riedasch for providing support and assistance in animal care and design of animal experiments. We are grateful for the Light Microscopy and Flow Cytometry Core Facilities of the German Cancer Research Center. We thank the Nikon Imaging Center of the Heidelberg University. We thank the Single-Cell Open Lab of the German Cancer Research Center. We thank the EM Core Facility of the Heidelberg University, the EM and Microscopy Core Facility of the German Cancer Research Center for their support.

## AUTHOR CONTRIBUTIONS

Supervision, V.V.; conceptualization, S.K.T., E.R. and V.V.; methodology, S.K.T., E.R., C.P.B., J.S., M.C.S., R.P.L., N.W. and V.V. ; project administration, S.K.T., E.R., V.V.; investigation, S.K.T., E.R., C.P.B., J.S., J.W., M.C.S, N.L., M.C.P., A.F., N.D., R.L.P., N.W., O.A., A.H., N.S., J.C., B.B., J.G.S., G.V., M.S., F.S. and V.V.; formal analysis, S.K.T., E.R., C.P.B., J.S., J.W., M.C.S., N.L., M.C.P., A.F., N.D., R.L.P., N.S., B.B., J.G.S., M.S., F.S. and V.V.; resources, O.A., J.C., G.V., K.F.N., C.A., B.S., D.H.H, V.V.; visualization, S.K.T., E.R., C.P.B., J.S., J.W., M.C.S, N.L., M.C.P., R.L.P., N.W., B.B., J.G.S., M.O.B. and V.V.; writing – original draft, V.V.; writing – review and editing, S.K.T., E.R., C.P.B., J.S, J.B., M.C.S, N.L., M.C.P., R.L.P., N.W., A.H., N.S., B.B., J.G.S., G.V., M.S., K.F.N., M.O.B., C.A., D.H.H, J.S.R., and V.V.; funding acquisition, V.V.

---

**Figure S1.**
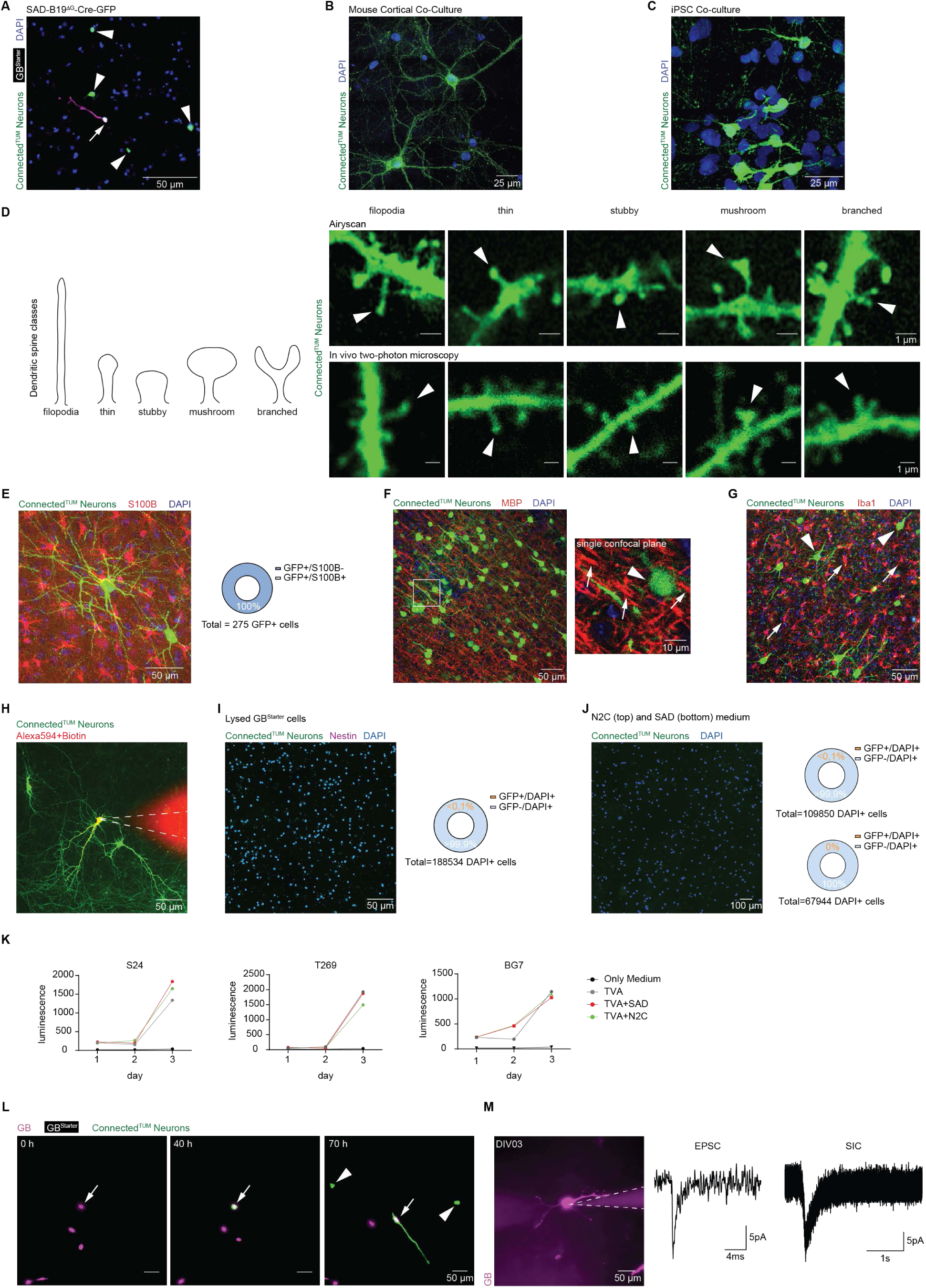
, Specific labeling of connected^TUM^ neurons with rabies-based tracing, related to Figure 1: (A) Probability map of SAD-B19^ΔG^-Cre-GFP and TVA-oG-mCherry positive S24 GB^Starter^ cell (arrow) with SAD-B19^ΔG^-Cre-GFP infected connected^TUM^ neurons (arrowheads) in co-culture. (B) Expansion microscopy of mouse cortical co-culture showing connected^TUM^ neurons in green (CVS-N2c^ΔG^-eGFP(EnvA), patient-derived model S24). (C) Expansion microscopy probability map of iPSC neuron co-culture showing connected^TUM^ neurons in green (CVS-N2c^ΔG^-eGFP(EnvA), patient-derived model S24). (D) Scheme illustrating dendritic spine classes (left). Dendritic spine classes can be distinguished in connected^TUM^ neurons (green) in PDX model S24 as shown with *ex vivo* Airyscan imaging (SAD-B19^ΔG^-eGFP(EnvA), right, top) and with *in vivo* two-photon imaging (CVS-N2c^ΔG^-eGFP(EnvA), right, bottom). Arrowheads point to dendritic spines matching the respective class. (E) *Ex vivo* maximum intensity projection of confocal microscopy from connected^TUM^ neurons (SAD-B19^ΔG^-eGFP(EnvA), green) and non-infected, S100B-positive astrocytes (red) in PDX model S24. All connected^TUM^ cells did not show S100B signal (n = 275 cells in n = 9 patient-derived models). (F) *Ex vivo* confocal maximum intensity projection showing connected^TUM^ neurons (CVS-N2c^ΔG^-eGFP(EnvA), green) and non-infected, MBP-positive oligodendrocytes (red) in PDX model S24 (left). Zoom-in on a single confocal plane showing no overlap between connected^TUM^ neurons (arrowhead) and MBP (arrows, right). (G) *Ex vivo* maximum intensity projection of confocal microscopy showing connected^TUM^ neurons (CVS-N2c^ΔG^-eGFP(EnvA), green, arrowheads) and non-infected, Iba1-positive microglia (red, arrows) in PDX model S24. (H) Representative image of whole-cell patch clamp of connected^TUM^ neuron (CVS-N2c^ΔG^-eGFP(EnvA)), patch pipette (dashed line) filled with Alexa 594 and Neurobiotin (patient-derived model S24). (I) Control experiment showing that rabies transfection from released particles from dead S24 GB^Starter^ cells (left). Less than 0.1% of cells in co-culture are infected. (n = 18 cells in n = 188534 cells total). (J) Culture medium from rabies-infected co-cultures on neuronal cultures (left). Quantifications show less than 0.1% of cells in culture are rabies-infected for both cultures infected with the CVS-N2c^ΔG^-eGFP(EnvA) (top) and the SAD-B19^ΔG^-eGFP(EnvA) (bottom) strains (n = 2 cells in n = 109850 cells total for CVS-N2c^ΔG^, n = 0 cells in n = 67944 cells total for SAD-B19^ΔG^ in patient-derived model S24). (K) CellTiter-Glo assay of patient-derived models S24 (left), T269 (middle) and BG7 (right) comparing only medium control (black), only TVA-oG-mCherry transduced cells (grey), SAD-B19^ΔG^-eGFP(EnvA) rabies-infected and TVA-oG-mCherry cells (red), and CVS-N2c^ΔG^-eGFP(EnvA) rabies-infected and TVA-oG-mCherry cells (green) (n = 2 replicates per model). (L) Time-lapse imaging showing a probability map of tumor cell dynamics immediately after seeding on co-cultures. Arrows pointing to newly infected GB^Starter^ cell, arrowheads indicating newly infected connected^TUM^ neurons (CVS-N2c^ΔG^-eGFP(EnvA)). (M) Image of S24 GB cell in whole-cell patch clamp recording 3 days after seeding onto neuronal cultures (left) and exemplary EPSC and SIC traces. Image was processed using denoised.ai.

**Figure S2.**
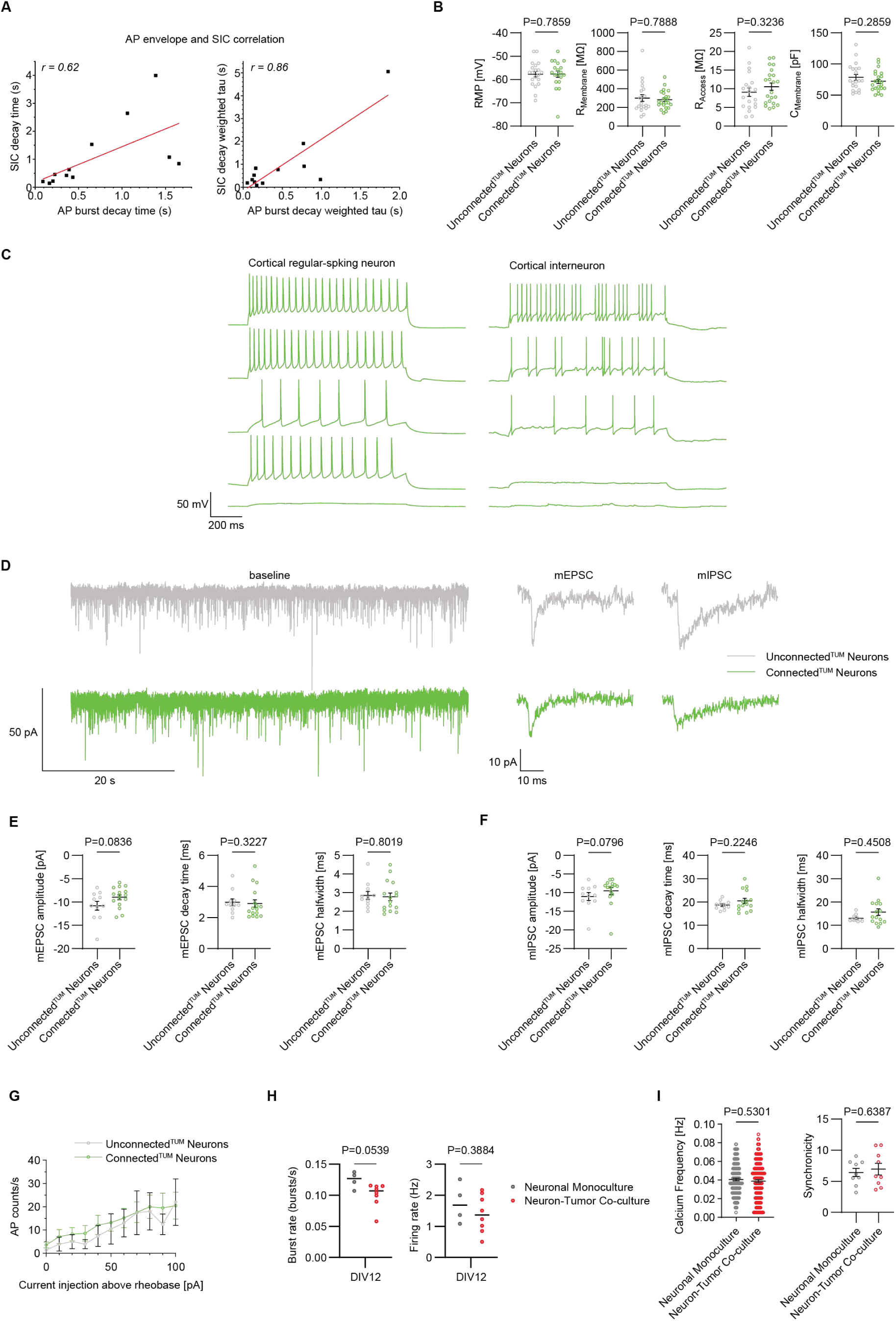
, Electrophysiological properties of connected^TUM^ and unconnected^TUM^ neurons, related to Figure 2: (A) Connected^TUM^ AP envelope and GB^Starter^ SIC current correlation, showing correlation between decay times (left, Pearson’s r = 0.62, ANOVA F (df) = 6.4 (11), p = 0.0301) and decay weighted tau (right, Pearson’s r = 0.86, ANOVA F (df) = 29.6 (11), p = 0.00029) (n = 12 pairs). (B) Passive membrane properties in currents in unconnected^TUM^ (n = 20) and connected^TUM^ (n = 22) cortical neurons: Resting membrane potential (RMP, Mann-Whitney test), membrane resistance (R_Membrane_, unpaired t-test), access resistance (R_Access_, unpaired t-test) and membrane capacitance (C_Membrane_, unpaired t-test). (C) Representative whole-cell current-clamp recordings of action potential firing in connected^TUM^ cortical regular- and intermittent-spiking neurons after 10, 50, 100, 150 and 200 pA current step injection. (D) Representative whole-cell voltage-clamp recordings of miniature post-synaptic currents in unconnected^TUM^ and connected^TUM^ cortical neurons with representative single mEPSC and mIPSC examples (right). (E) Post-synaptic mEPSC properties in unconnected^TUM^ (n = 11) and connected^TUM^ (n = 16) cortical neurons: mEPSC amplitude (unpaired t-test), decay time (Mann-Whitney test) and half-width (unpaired t-test). (F) Post-synaptic mIPSC properties in unconnected^TUM^ (n = 11) and connected^TUM^ (n = 16) cortical neurons: mEPSC amplitude (unpaired t-test), decay time (unpaired t-test) and half-width (Mann-Whitney test). (G) Burst rate in bursts/s (left) and firing rate in Hz (right) of cultures without tumor cells (grey) and cultures with GB (red) (n = 4 monocultures and 8 co-cultures, unpaired t-test). (H) Calcium transient frequency (left) and synchronicity (right) of neuronal monoculture and cultures with seeded GB cells (n = 157 (monoculture) and 160 (co-culture) cells in 9 regions of interest, Mann-Whitney test (frequency) and unpaired t-test (synchronicity)). (I) Input-output relationship between the current injected (relative to the rheobase current) and the number of action potentials generated over 1 s in connected^TUM^ and unconnected^TUM^ intermittent-spiking neurons (n = 8).

**Figure S3.**
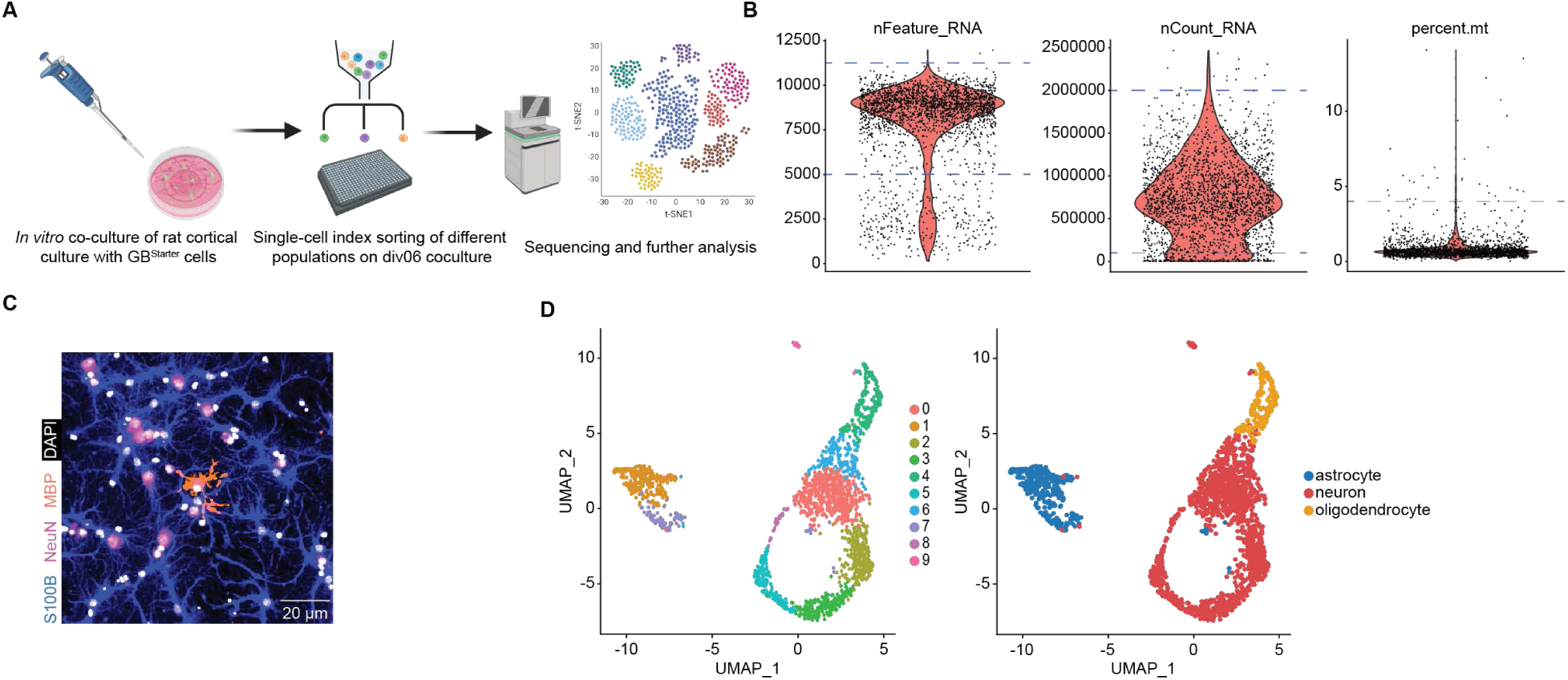
, Single-cell RNA sequencing of connected^TUM^ and unconnected^TUM^ neurons, related to Figure 2: (A) Schematic workflow of the FACS sorting and single-cell RNA sequencing of co-cultures. (B) Quality control of sequenced co-cultures. Blue dashed lines indicate filtering cut-offs. (C) Representative image showing the different microenvironmental cell types in co-culture. The main portion of the cells found are S100B-positive astrocytes (blue), NeuN-positive neurons (magenta) and MBP-positive oligodendrocytes (orange). (D) UMAP plots of the sequenced co-cultures after quality control showing the clustering (left) and the cell type annotation (right) of the different microenvironmental cell types in co-culture (n = 1958 cells).

**Figure S4.**
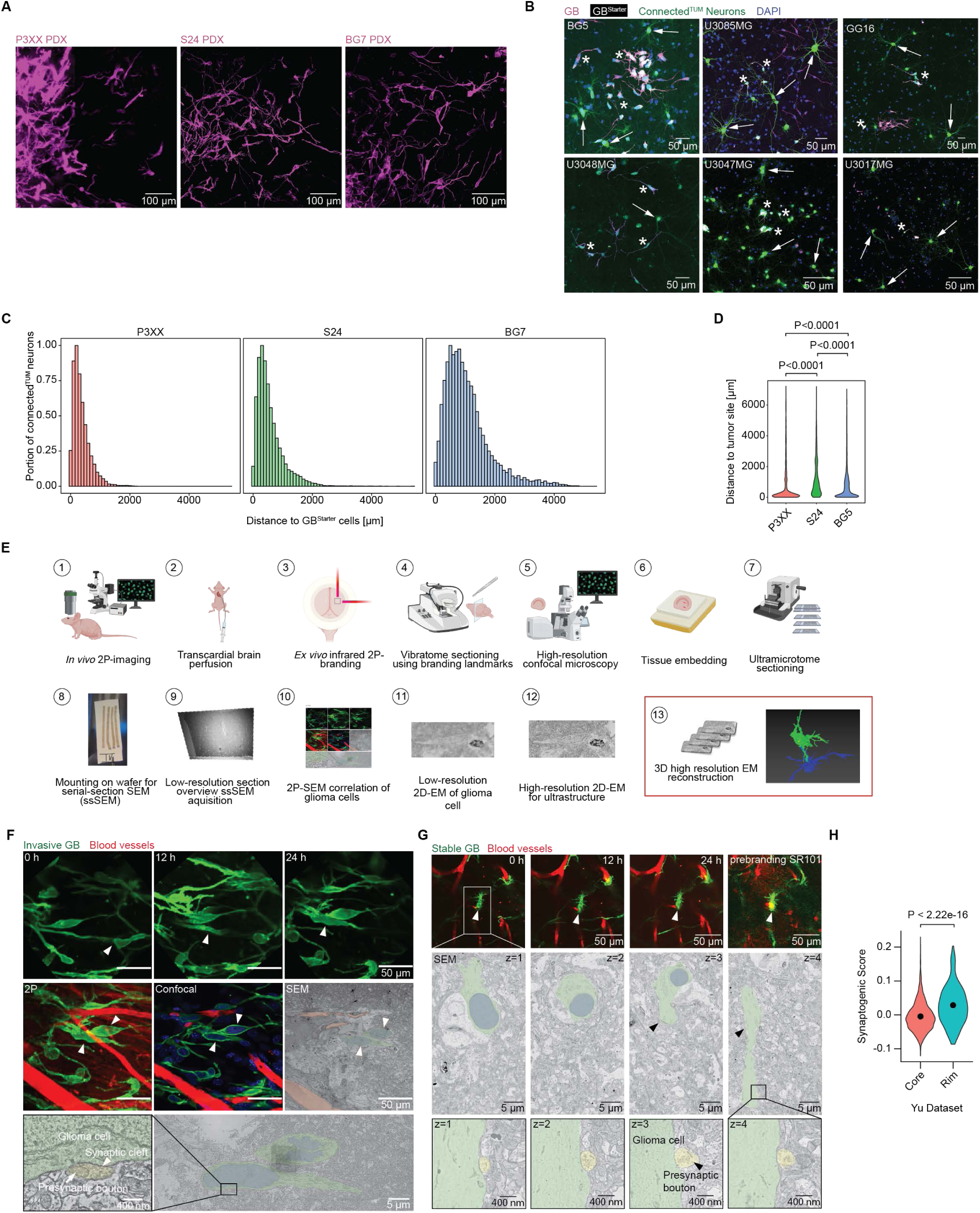
, Neuron-to-glioma synapses across functional glioblastoma cell states, related to Figure 3: (A) *In vivo* two-photon microscopy images of 3 different PDX models P3XX (left), S24 (middle) and BG7 (right). Images were processed with denoise.ai. (B) Neuronal connectome of different patient-derived glioblastoma models in co-culture. Representative images showing GB^Starter^ cells (white, asterisks) and their connected^TUM^ neurons (green, arrows) (CVS-N2c^ΔG^-eGFP(EnvA) for BG5, GG16, U3048MG, SAD-B19^ΔG^-eGFP(EnvA) for U3085MG, U3047MG, U3017MG, green). (C) Histogram showing the portion of connected^TUM^ neurons in relation to the distance to GB^Starter^ cells for patient-derived models P3XX (left), S24 (middle) and BG7 (right) in co-culture (n = 30219 (S24), n = 17726 (P3XX), n = 10877 (BG7) connected^TUM^ neurons in n = 3 biological replicates). (D) Distance of connected^TUM^ neurons to main tumor site in three PDX models *in vivo* (n = 17726 (P3XX), n = 30219 (S24), n = 10877 (BG7) cells, one-way ANOVA test). (E) *In vivo* correlative light and serial section scanning electron microscopy (CLEM) workflow. (F) CLEM of an invasive GB cell with 24h *in vivo* two-photon imaging prior to perfusion and subsequent electron microscopy revealing a neuron-to-glioma synapse with presynaptic bouton and synaptic cleft. Glioblastoma cell (green overlay), its nucleus (blue overlay), presynaptic bouton (yellow overlay) and blood vessels (red overlay). (G) CLEM of a stable GB cell with 24h *in vivo* two-photon imaging prior to perfusion with SR101-signal in the cell. Serial sectioning scanning electron microscopy of the glioblastoma cell (green overlay) and its nucleus (blue overlay). Zoom-in on a neuron-to-glioma synapse in consecutive z-layers with a presynaptic bouton (yellow overlay). (H) Synaptogenic score in rim compared to core glioblastoma regions in the Yu dataset^63^ (n = 2795 cells, Mann-Whitney test).

**Figure S5.**
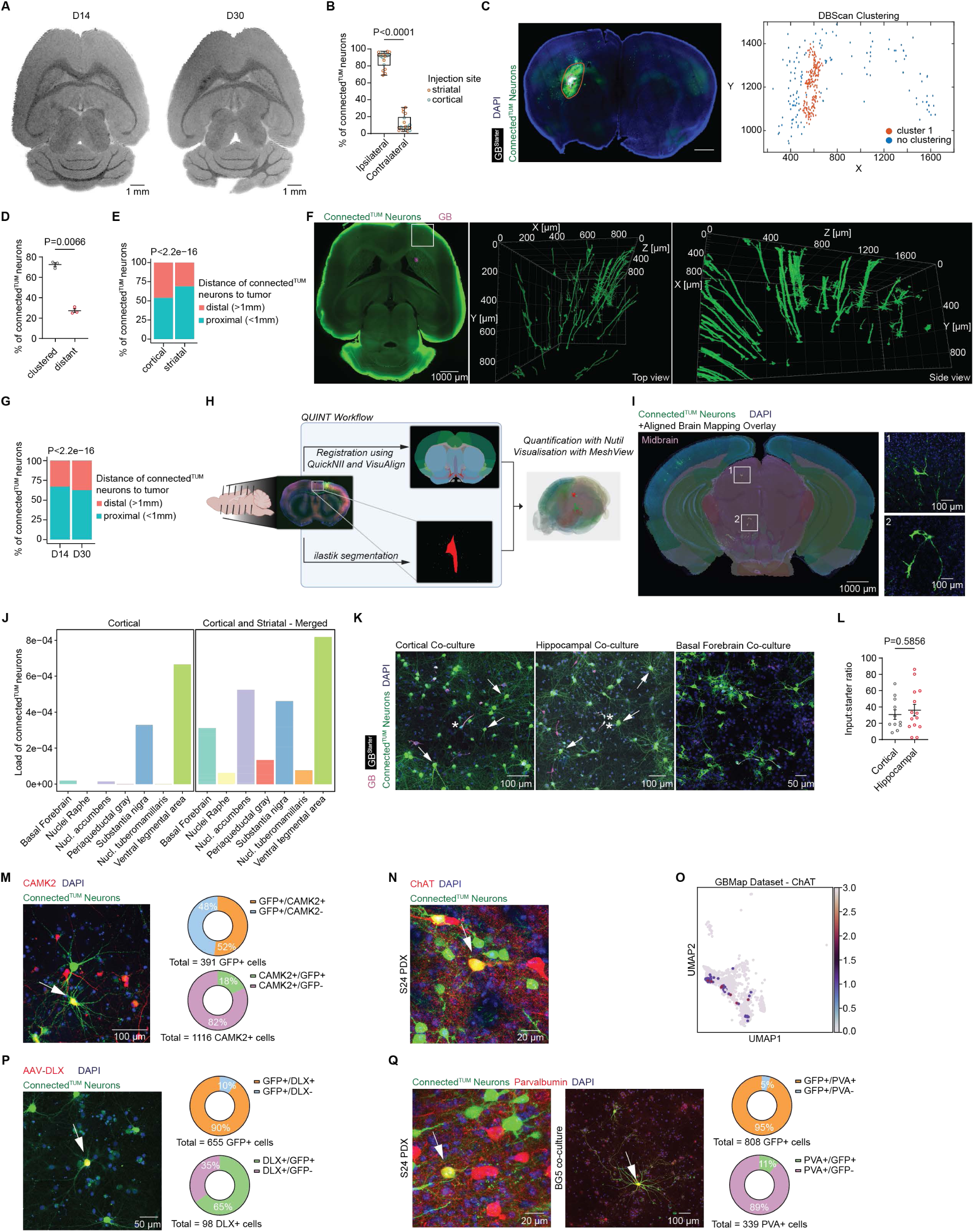
, Brain atlas mapping of connected^TUM^ neurons, related to Figure 4: (A) MRI imaging of early stage tumors (D14 and D30) showing no T2-signal (PDX model BG5). (B) Comparison of the portion of connected^TUM^ neurons in the ipsilateral and contralateral hemispheres in relation to the tumor site in cortical and striatal tumors (n = 7 cortical and n = 11 striatal tumors from three PDX models (S24, BG5, P3XX), Mann-Whitney test). (C) Exemplary *ex vivo* brain slice image (left) showing S24 GB^Starter^ cells (white) and connected^TUM^ neurons (green). Dashed circle points to the tumor site. Orange circle indicating the area of the majority of connected^TUM^ neurons. Scale bar = 1 mm. DBScan clustering of the connected^TUM^ neurons from the image on the left (right). One big cluster around the tumor site is detected (cluster 1, orange). More distant connected^TUM^ neurons show no specific clustering (blue). (D) Proportion of connected^TUM^ neurons in clusters compared to distant connected^TUM^ neurons in co-culture (n = 3 samples from patient-derived models S24 and BG7). Clusters were determined with DBScan clustering. (E) Comparison of the portion of proximal and distal connected^TUM^ neurons in cortical compared to striatal tumors (n = 8839 connected^TUM^ neurons in n = 7 cortical tumors, n = 30528 connected^TUM^ neurons in n = 11 striatal tumors in three PDX models (S24, BG5, P3XX), Wilcoxon test). (F) Light-sheet microscopy of retrograde tracing of an early stage glioblastoma (PDX model BG5, D30 post tumor injection). Single plane image showing tumor (magenta) and connected^TUM^ neurons (CVS-N2c^ΔG^-eGFP(EnvA), green, left). 3D renderings showing zoom-in onto the connected^TUM^ neurons in the marked region on the right from two different perspectives. (G) Comparison of the portion of proximal and distal connected^TUM^ neurons 14 versus 30 days after tumor implantation (n = 26419 connected^TUM^ neurons in n = 11 D14 tumors, n = 12948 connected^TUM^ neurons in n = 7 D30 tumors in three PDX models (S24, BG5, P3XX), Wilcoxon test). (H) Scheme of the QUINT workflow for atlas mapping of brain sections (STAR Methods). (I) Overlay of fluorescence microscopy and brain atlas mapping around the midbrain region (PDX model S24, left). Zoom-in on connected^TUM^ neurons in the brainstem (SAD-B19^ΔG^-eGFP(EnvA), right). (J) Bar plot showing the load of connected^TUM^ neurons in various neuromodulatory circuits in cortical tumors (left) and in all analyzed samples (right) (n = 8839 connected^TUM^ neurons in n = 7 cortical tumors, n = 39367 connected^TUM^ neurons in n = 18 mice total from three PDX models (S24, BG5, P3XX)). (K) Representative confocal images of different co-culture models with neurons from different brain regions. Rat cortical co-culture (left), hippocampal co-culture (middle) and basal forebrain co-culture (right). Asterisks point to S24 GB^Starter^ cells, arrows showing exemplary connected^TUM^ neurons (CVS-N2c^ΔG^-eGFP(EnvA) in cortical and hippocampal culture, SAD-B19^ΔG^-eGFP(EnvA) in basal forebrain culture). (L) Input-to-starter ratio in hippocampal compared to cortical co-culture model (n = 11 samples for cortical, n = 14 samples for hippocampal cultures, unpaired t-test).) (M) Representative confocal image of connected^TUM^ neurons (SAD-B19^ΔG^-eGFP(EnvA), green) and CAMK2-positive neurons (red, left). Arrow points to a CAMK2-positive connected^TUM^ neuron (yellow). Analysis illustrating the portion of CAMK2-positive connected^TUM^ neurons compared to all connected^TUM^ neurons (above right, n = 391 connected^TUM^ neurons in 3 biological replicates). Analysis showing the portion of connected^TUM^ neurons compared to all CAMK2-positive neurons (below right, n = 1116 CAMK2-positive cells in 3 biological replicates). (N) Representative confocal image of connected^TUM^ neurons (SAD-B19^ΔG^-eGFP(EnvA), green) and ChAT-positive neurons (red). Arrow points to a ChAT-positive connected^TUM^ neuron (yellow). (O) UMAP plot of the neuronal cell subpopulation in the GBMap dataset^114^ showing ChAT expression (n = 6309 neurons). (P) Exemplary confocal image of connected^TUM^ neurons (CVS-N2c^ΔG^-eGFP(EnvA), green) in a co-culture infected with AAV-DLX virus infecting GABAergic interneurons (left)^132^ (red). Arrow points to a DLX-positive connected^TUM^ neuron. Analysis illustrating the portion of DLX-positive connected^TUM^ neurons compared to all connected^TUM^ neurons (above right, n = 655 connected^TUM^ neurons in n = 6 samples). Analysis showing the portion of connected^TUM^ neurons compared to all DLX-positive neurons (below right, n = 98 DLX-positive cells in n = 6 samples). (Q) Exemplary confocal images of connected^TUM^ neurons (SAD-B19^ΔG^-eGFP(EnvA), green) and Parvalbumin-expressing neurons (red) in a patient-derived xenograft model (left) and in co-culture (middle). Arrows point to Parvalbumin-positive connected^TUM^ neurons (yellow). Analysis illustrating the portion of Parvalbumin-positive connected^TUM^ neurons compared to all connected^TUM^ neurons (above right, n = 808 connected^TUM^ neurons in n = 4 biological replicates). Analysis showing the portion of connected^TUM^ neurons compared to all Parvalbumin-positive neurons (below right, n = 339 Parvalbumin-positive cells in n = 4 biological replicates).

**Figure S6.**
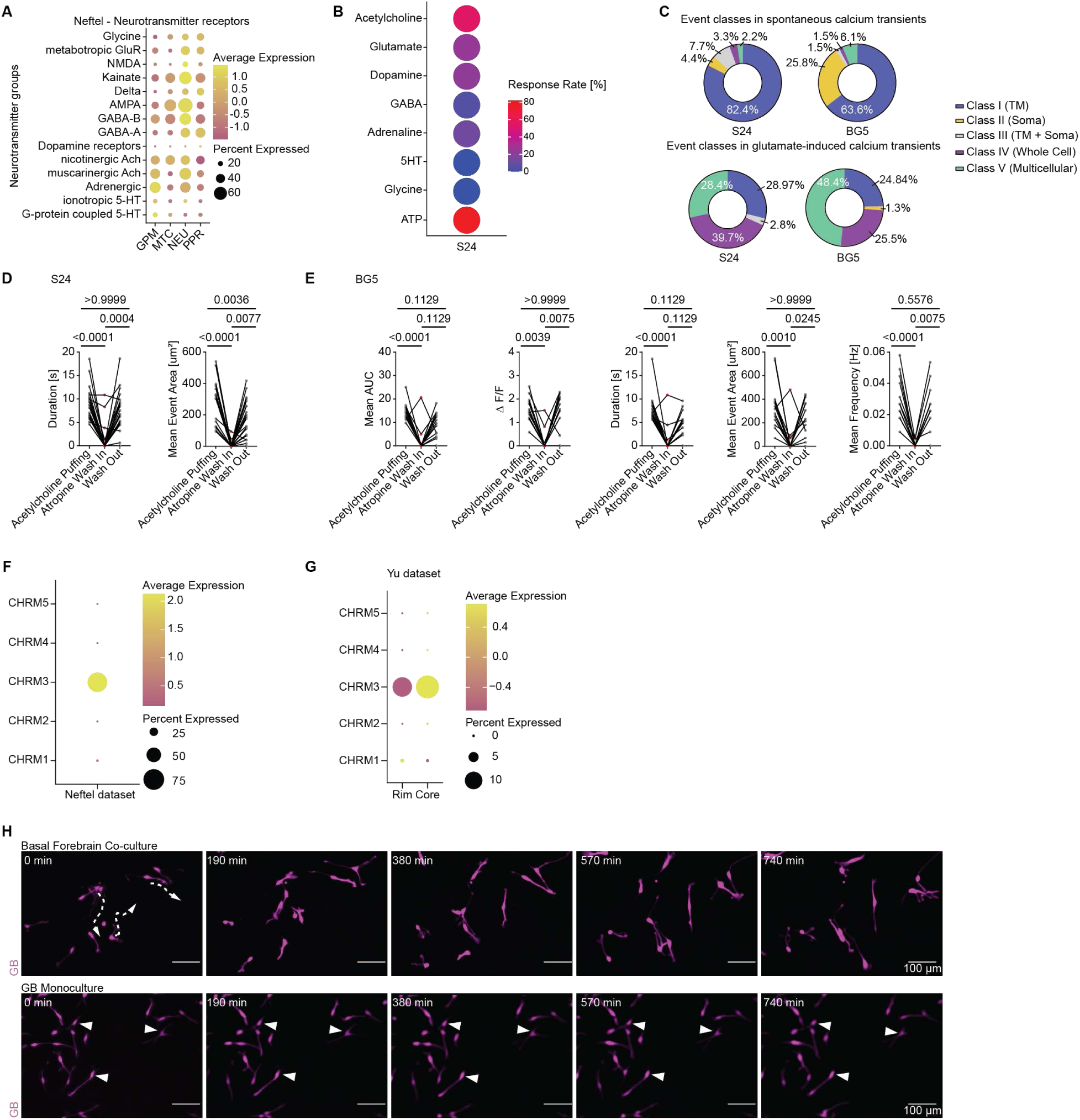
, Gene expression and functional neurotransmitter receptor analysis in glioblastoma, related to Figure 4: (A) Dot plot showing the gene expression module scores of various neurotransmitter receptor groups of different glioblastoma pathway-based cell states^7^ in the Neftel dataset^5^ (n = 7929 cells). (B) Dot plot indicating the calcium transient response rate to stimulation with different neurotransmitters in patient-derived model S24 (n = 56 cells from 5 independent experiments). (C) Calcium transient event classes in S24 and BG5 patient-derived models under spontaneous baseline conditions (above) and induced by glutamate puffing (below) (n = 91 events in n = 29 cells (S24), n = 65 events in n = 56 cells (BG5) for spontaneous events; n = 176 events in n = 77 cells (S24) and n = 157 events in n = 66 cells (BG5) in 12 (S24) and 6 (BG5) independent experiments). (D) Duration (left) and mean event area (right) of calcium transients in response to acetylcholine puffing, inhibition of transients by atropine and wash out in S24 glioblastoma cells (n = 22 cells from 2 independent experiments, Friedman test). (E) Mean area under curve, ΔF over F, duration, event area und frequency (from left to right) of calcium transients in BG5 glioblastoma cells responding to acetylcholine puffing, inhibition of transients by atropine and wash out (n = 14 cells from one experiment, Friedman test). (F) Dot plot showing the expression of muscarinergic acetylcholine receptors in glioblastoma cells in the Neftel dataset^5^ (n = 7929 cells). (G) Dot plot showing the gene expression of muscarinergic acetylcholine receptor subunits in glioblastoma cells split by rim versus core in Yu dataset^63^ (n = 2795 cells). (H) *In vitro* live cell time-lapse images of glioblastoma cells in a co-culture of tumor cells and basal forebrain neurons (top) compared to a monoculture of only glioblastoma cells (bottom). Arrows with dashed lines indicating movement of invasive cells, arrows pointing to stable cells. Images were processed with denoise.ai.

**Figure S7.**
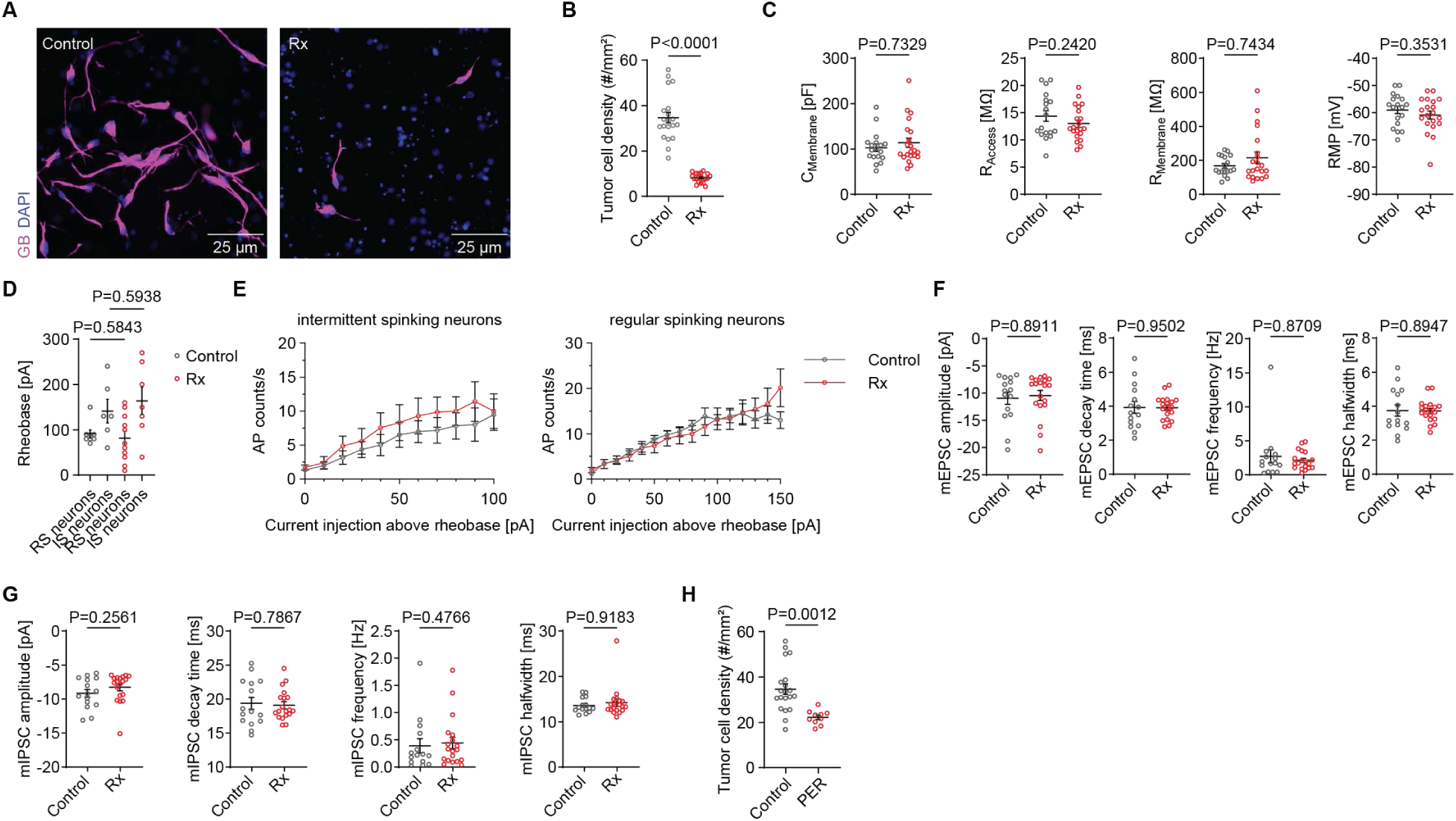
, Radiotherapy-induced effects on glioblastoma cells and connected^TUM^ neurons, related to Figure 6: (A) Representative images of tumor regions in control (left) compared to radiotherapy-treated (right) conditions. (B) Tumor cell density in cell count per mm^2^ under control conditions versus after irradiation (n = 20 control versus 20 irradiated samples, unpaired t-test) (C) Passive membrane properties of connected^TUM^ cortical neurons under control condition and after radiotherapy: Membrane capacitance (C_Membrane_), access resistance (R_Access_), membrane resistance (R_Membrane_) and resting membrane potential (RMP) (n = 18 control and n = 20 neurons after radiotherapy, Mann-Whitney test for C_Membrane_ and R_Membrane_, unpaired t-test for RMP and R_Access_). (D) Neuronal rheobase of connected^TUM^ cortical neurons under control condition and after radiotherapy split by neuronal firing type (n = 8 regular-spiking control neurons, n = 11 regular-spiking neurons after radiotherapy, Mann-Whitney test, n = 6 intermittent-spiking control neurons, n = 7 intermittent-spiking neurons after radiotherapy, unpaired t-test) (E) Input-output relationship between the current injected relative to the rheobase current and the number of action potentials generated over 1 s in connected intermittent-spiking and regular-spiking neurons. (F) Post-synaptic mEPSC properties of connected^TUM^ neurons under control conditions (n = 15) and after radiotherapy (n = 19) (Mann-Whitney test for mEPSC amplitude and frequency; unpaired t-test for mEPSC half-width and decay time). (G) Post-synaptic mIPSC properties of connected^TUM^ neurons under control conditions (n = 15) and after radiotherapy (n = 19) (Mann-Whitney test for mIPSC amplitude, frequency and half-width; unpaired t-test for mIPSC decay time). (H) Tumor cell density in cell count per mm^2^ under control conditions compared to after perampanel treatment (n = 20 control versus n = 10 perampanel-treated samples, unpaired t-test).

